# An integrated workflow for crosslinking mass spectrometry

**DOI:** 10.1101/355396

**Authors:** Marta L. Mendes, Lutz Fischer, Zhuo A. Chen, Marta Barbon, Francis J. O’Reilly, Sven Giese, Michael Bohlke-Schneider, Adam Belsom, Therese Dau, Colin W. Combe, Martin Graham, Markus R. Eisele, Wolfgang Baumeister, Christian Speck, Juri Rappsilber

## Abstract

We present a concise workflow to enhance the mass spectrometric detection of crosslinked peptides by introducing sequential digestion and the crosslink identification software Xi. Sequential digestion enhances peptide detection by selective shortening of long tryptic peptides. We demonstrate our simple 12-fraction protocol for crosslinked multi-protein complexes and cell lysates, quantitative analysis, and high-density crosslinking, without requiring specific crosslinker features. This overall approach reveals dynamic protein-protein interaction sites, which are accessible, have fundamental functional relevance and are therefore ideally suited for the development of small molecule inhibitors.

---

Crosslinking mass spectrometry (CLMS) has become a standard tool for the topological analysis of multi-protein complexes and has begun delivering high-density information on protein structures, insights into structural changes and the wiring of interaction networks *in situ*^1^. The technological development currently focuses on enrichment strategies for crosslinked peptides and mass spectrometric data acquisition^2–4^, including newly designed crosslinkers^5^. MS2-cleavable crosslinkers in particular, have celebrated recent successes for the analysis of protein complexes^6^ or complex mixtures^7,8^. Functionalized crosslinkers typically form part of an entire workflow including analysis software tailored to the data formats obtained, such as XLinkX^7^ but, importantly, there are also flexible general software that are not paired with distinct crosslinker properties, such as pLink^9^, StavroX^10^ and Kojak^11^.

The focus on bespoke crosslinkers has left general steps of sample preparation, such as protein digestion, with less attention. Tryptic digestion generates crosslinked peptides of considerable size; a quality that has been exploited with their enrichment by SEC^12^, but one that poses as a potential problem regarding their detection. Replacing trypsin with proteases such as GluC, AspN and chymotrypsin does not change peptide size distributions fundamentally^13^. We reasoned that sequential digestion could reduce the size of large tryptic peptides and offer access to sequence space that otherwise would remain undetected. We therefore followed trypsin digestion with subsequent digestion by alternative proteases and developed Xi, a database search engine, allowing the search of multiple datasets resulting from the application of our protocol. This novel approach expands the detectable structure space in proteins, allowing it to capture dynamic regions in protein complexes that are mechanistically important and druggable, however that hitherto have remained undisclosed by cryo-EM.

We first tested this workflow on a standard mix of seven Bis[sulfosuccinimidyl] suberate (BS^3^) crosslinked proteins (Catalase, Myoglobin, Cytochrome C, Lysozyme, Creatine Kinase, HSA and Conalbumin). Importantly, their structures are known and hence offer an independent assessment of false identifications. Four digestion conditions, each giving three SEC fractions, resulted in a total of 12 acquisitions, which is the protocol applied to all subsequent analyses presented here (Fig. 1a). The results of this protocol for our standard proteins was compared to a parallel digestion using the same four enzymes and using trypsin alone in four replica, maintaining the analytical effort comparable in all three cases (SEC fractionation, 12 injections). Sequential digestion produced the best results when compared to replica analyses and parallel digestion (Fig. 1b,c, Supplementary Fig. 1a, Supplementary Data 1). Before assessing if this improvement translated into a gain of information in biological applications, we investigated the origin of the added data.

**Figure 1:**
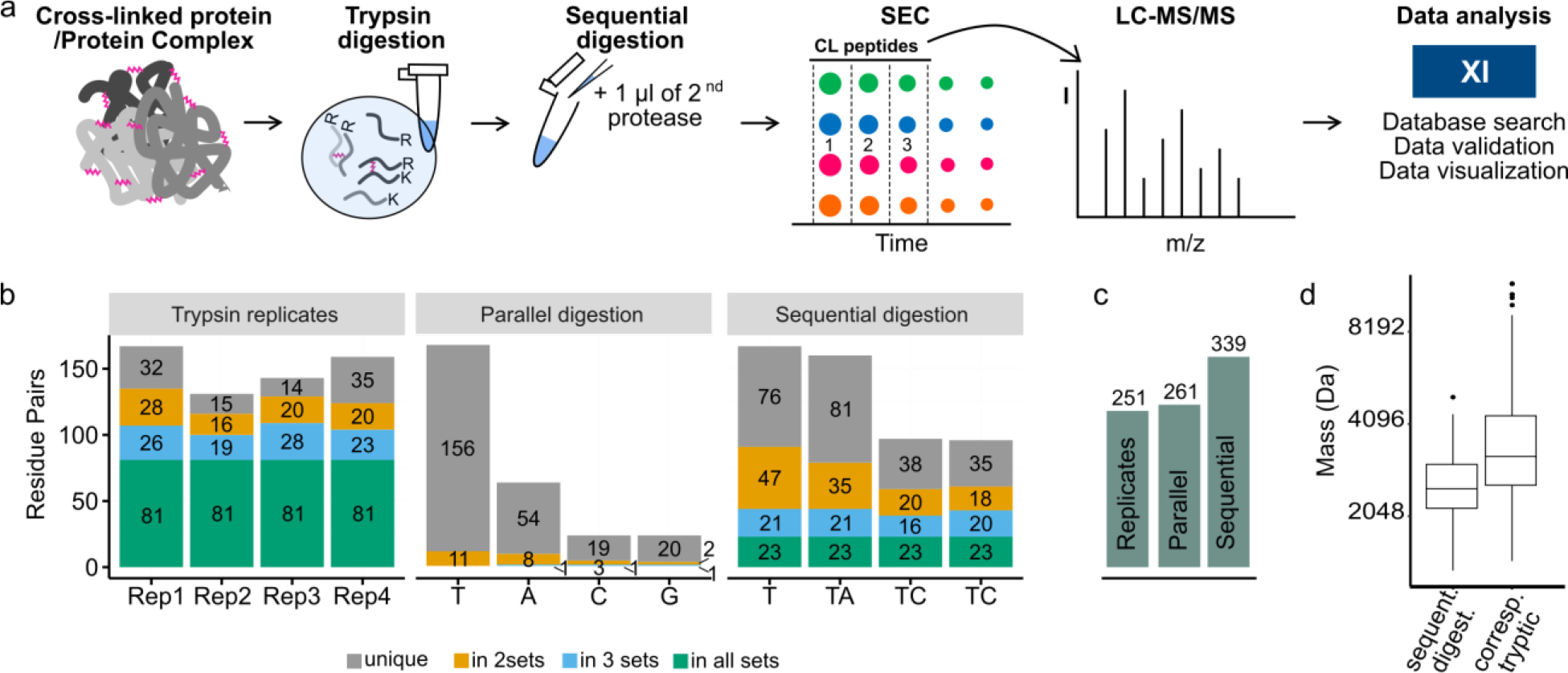
*Sequential digestion strategy and its impact on different sample complexities, crosslinker chemistry and quantitation. (a) Sequential digestion workflow. Proteins or protein complexes are crosslinked and digested with trypsin. After splitting the sample into four aliquots, one remains single digested with trypsin (T) while the others are sequentially digested with either AspN (A), chymotrypsin (C) or GluC (G). Samples are enriched by SEC and the three high-MW fractions are analysed by LC-MS, submitted to Xi search and xiFDR analysis. (b) Overlap of the number of residue pairs for sequential digestion and the control experiments composed by an experiment using trypsin alone in four replicates and individual digestions with trypsin, AspN, chymotrypsin and GluC. A trypsin four replicate experiment shows a large overlap of the four datasets with little gain. Parallel digestions with trypsin, AspN, chymotrypsin and GluC demonstrates high complementarity but moderate gains over trypsin. Sequential digestion shows low overlap between the four datasets and the largest gain in unique residue pairs. (c) Gains of repeated analysis (trypsin only), parallel digestions and sequential digestion. (d) Crosslinked peptides obtained by sequential digestion are smaller than their corresponding tryptic peptides*.

Indeed, sequential digestion led to smaller peptides than trypsin alone (Fig. 1d, Supplementary Fig. 2f) and moved the mass distribution of theoretical crosslinkable peptides more into the mass range typically detected by our instrument (Supplementary Fig. 2f). For short peptides we noticed a protection effect, based on the number of peptides containing missed cleavage sites and on the number of missed cleavage sites relative to peptide length (Supplementary Fig. 3). This agrees with reports that serine proteases lose efficiency as peptides shorten^14,15^. Although AspN is a metalloprotease, it showed a similar loss of efficiency for short peptides. Notably, we observed a bias towards maintaining tryptic C-termini. Crosslinked peptides with two tryptic C-termini are more frequently identified while those with C-termini generated by the second protease are less frequent than expected, relying on N-termini as internal reference (Supplementary Fig. 1c). This identification bias is consistent with better fragmentation behaviour of peptides with basic C-termini^16^ and testifies to the importance of trypsin as part of the protocol.

**Figure 2:**
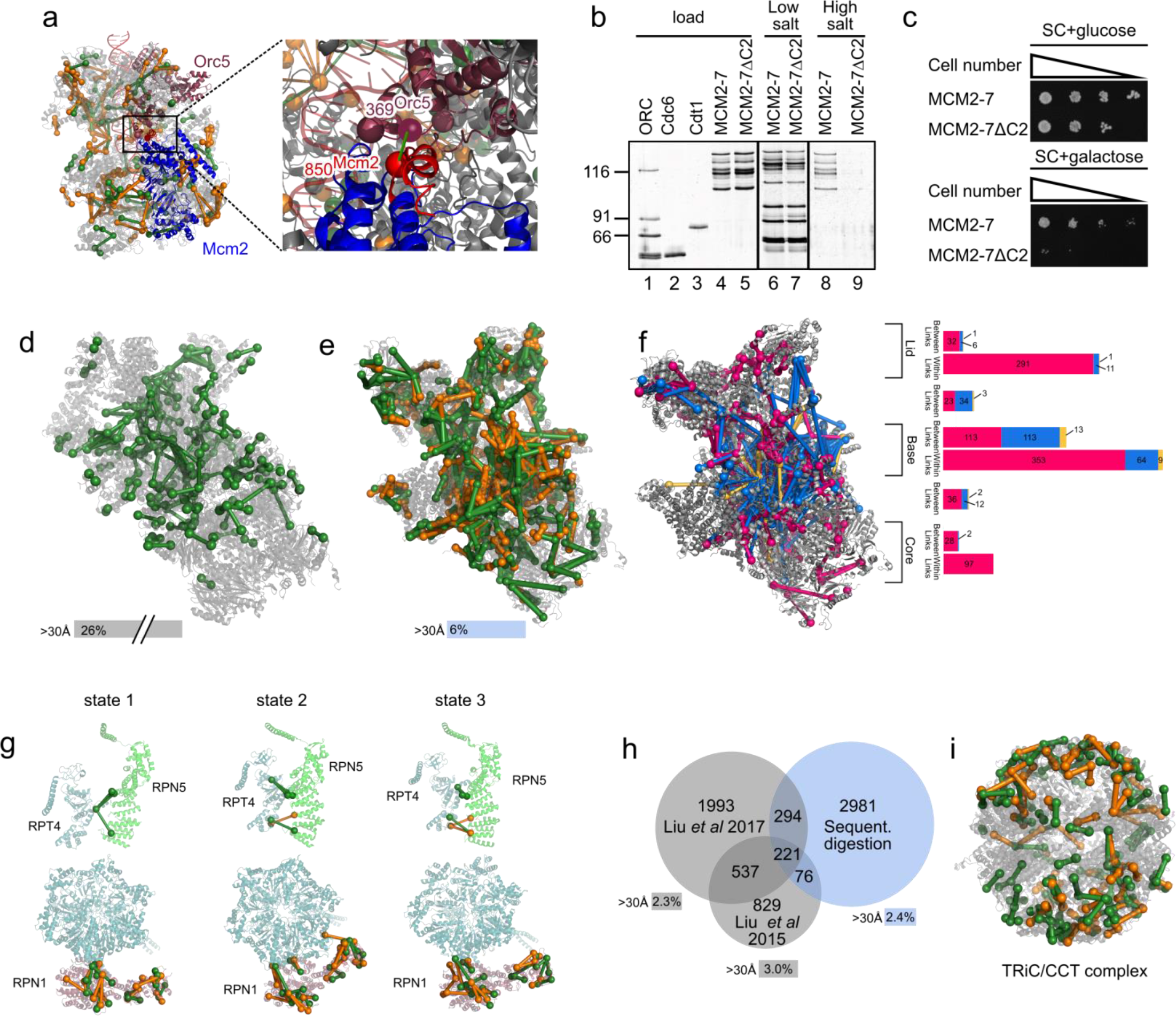
*Sequential digestion of the affinity-purified complexes OCCM* (S. cer.*) and 26S proteasome (*S. cer.*) and the cytosol of human K562 cells. Residue pairs observed in tryptic (green) and non-tryptic (orange) peptides. (a) Unique residue pairs mapped to the OCCM complex (PDB|5udb) and the key link Mcm2 – Orc5 (Mcm2-850-Orc5-369). (b) The in vitro helicase loading assay demonstrates that an Mcm2 C-terminal deletion mutant supports complex assembly (lanes 6 and 7) and blocks formation of the final helicase loading product (lanes 8 and 9). (c) Overexpression analysis of Mcm2-7ΔC2 shows that this mutant causes dominant lethality, indicating that the C-terminus of Mcm2 is essential in cell survival. (d) Unique residue pairs obtained by Wang* et al. *for the human 26S proteasome (PDB|5GJR). (e) Unique residue pairs obtained by sequential digestion obtained for the S. cerevisiae 26S proteasome (PDB|4CR2). Sequential digestion returned the highest number of residue pairs so far identified by CLMS for the 26S proteasome. (f) Long distance (blue) and within distance (pink) between residue pairs were mapped into one of the states of the proteasome (4cr2) showing the accumulation of those into the base of the complex. Residue pairs satisfying other states are represented in yellow. The bar plot shows the distribution of all residue pairs in the complex showing that long distance links locate mainly in the base. (g) Unique residue pairs were mapped into the 3 states described by Unverdorben et al. showing that different conformations can be distinguished by using sequential digestion. (h) Comparison of sequential digestion for complex mixtures with previous studies. Sequential digestion returns a higher number of residue pairs with low overlap to published datasets, showing the complementarity of the different approaches. Over-length links were determined only within proteins and are comparable for all datasets. (i) Residue pairs for the TRiC/CCT complex were mapped into the crystal structure and support the rearrangement of the complex reported by Leitner et al.^1^(PDB|4V94).*

**Figure 3:**
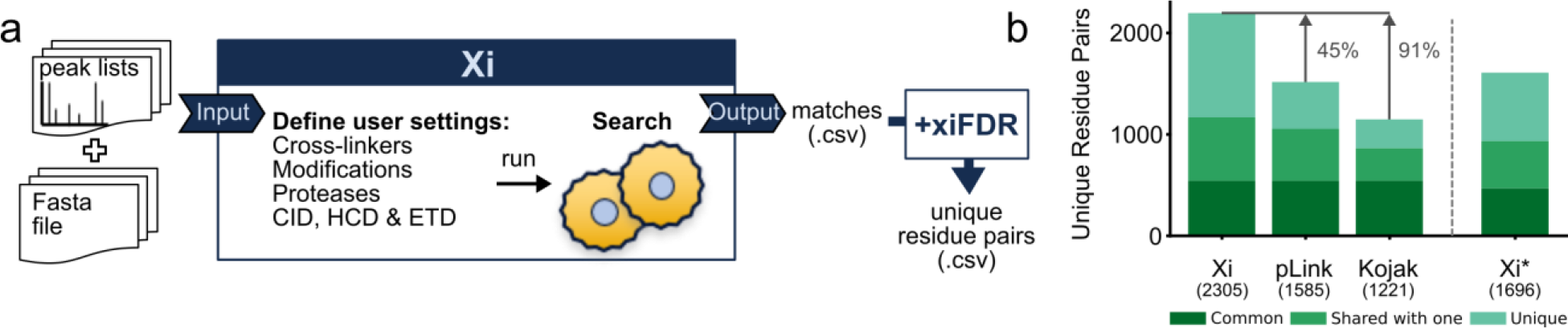
*Xi search engine. (a) Xi is an open source search engine that takes a peak list as input. Users can define any type of crosslinker, modification, digestion, and fragmentation method. The output is a list of matches in .csv format. We use xiFDR to filter results to the desired confidence level. (b) Xi(+xiFDR), pLink and Kojak(+PeptideProphet) comparison at 5% residue-pair FDR. The same trypsin dataset of the 26S proteasome was searched with all three software packages. Xi was run twice – once giving same likelihood for matching Lysine, Serine, Threonine and Tyrosine (Xi), as is the case for Kojak and pLink, and once giving priority to Lysine (Xi*)*.

We then tested the sequential digestion approach on samples of increasing complexity ranging from single proteins, UGGT and C3b, to the OCCM DNA replication complex (1.1 MDa), the 26S proteasome (2.5 MDa) and high-molecular weight fractions of human cytosol. A quantitative experiment was performed to assess the efficiency of sequential digestion combined with the QCLMS workflow^17^. Additionally, we tested the approach using two different crosslinkers, the homobifunctional crosslinker BS^3^ and the heterobifunctional, photoactivatable crosslinker sulfosuccinimidyl 4,4’-azipentanoate (SDA).

UGGT was one of the data-assisted *de novo* folding targets of CASP12 for which we contributed data in the form of 433 unique residue pairs obtained at a 5% FDR (http://predictioncenter.org/download_area/CASP12/extra_experiments/) (Supplementary Fig. 4a) using SDA as crosslinker and 26 LC-MS runs^18^. Using sequential digestion, we now identified 1523 unique residue pairs in only 12 runs (Supplementary Fig. 4b, c, Supplementary Data 1 and 2). With 5% long-distance links (> 20 Å) when mapped onto the structure released by CASP organizers (Supplementary Fig. 4d) the 3-fold increase in observed links comes at uncompromised reliability. Consequently, the sequential digestion protocol improves high-density CLMS by a clear increase in the number of residue pairs while simultaneously reducing the analytical effort needed to detect these.

We next combined quantitative CLMS (QCLMS)^19–21^ with our workflow (Supplementary Fig. 5a) to investigate the dimerization of C3b. Thioester-mediated dimerization of C3b is a key process of the human complement response enhancing the efficiency of C5 convertase formation which ultimately leads to clearance of pathogens from human blood^22–24^. However, the structure of this dimer is currently unclear. The reactive thioester could result in a random orientation of the two C3b molecules in a dimer. Alternatively, auxiliary factors or self-organisation properties of C3b could mediate a preferred orientation. We here investigate C3b alone and find it to form dimers in the absence of active thioester and auxiliary proteins. We quantified 293 unique crosslinks, about three times more than with trypsin alone (99) (Supplementary Fig. 5c, Supplementary Data 1 and 2) which lends robust support to a bottom-to-bottom orientation (Supplementary Fig. 5d). This suggests non-covalent interactions between C3b molecules lead to a preferred dimer orientation which implies that a thioester bridged dimer would follow this arrangement. Non-covalent interactions thus self-organise C3b into a productive dimer as this arrangement is compatible with the subsequent molecular events of the complement cascade by allowing unhindered binding of factor B at the top of C3b.

Turning our attention to protein complexes we investigated the OCCM complex, a helicase-loading intermediate formed during the initiation of DNA replication. Recently a 3.9-Å structure of *Saccharomyces cerevisiae* OCCM on DNA was obtained by cryo-electron microscopy (cryo-EM), supported by CLMS.^25^ We identified 682 residues pairs from the same sample analysed before, with large contribution from sequential digestion (Fig 2a, Supplementary Fig. 6a,b, Supplementary Data 1 and 2). Interactions observed now include known Cdt1-Mcm2 and Mcm6 but also Mcm2-Orc5 interaction (Mcm2-850-Orc5-369). These led us to delete the C-terminal 20 aa of Mcm2 (848-868) (Fig. 2C lane 5) and analyse its biological relevance in a well-established *in vitro* helicase loading assay, which recapitulates the *in vivo* process^26^. The deletion mutant did not affect ORC, Cdc6, Cdt1 and origin DNA dependent complex assembly under low salt conditions (Fig. 2b, lanes 6 and 7), but severely impaired the formation of the high salt stable double-hexamer (Fig. 2b, lanes 8 and 9), the final product of the helicase loading reaction. This is an exciting result, as it highlights a novel and essential role for Mcm2 C-terminus in a late step of MCM2-7 double-hexamer formation, a process that is only poorly understood. Moreover, the CLMS data show that the Mcm2 C-terminus is involved in a network of interactions with flexible domains of Orc6, Orc2 and Mcm5, which could represent an ideal target for development of inhibitors with potential as anti-cancer therapy^27^, as dynamic interactions have improved druggability characteristics over stable protein interactions^28,29^. Indeed, expressing Mcm2-7ΔC2 causes dominant lethality (Fig. 2c). The ability of CLMS data to complete the cryo-EM structure of the OCCM complex by dynamic contacts proved here essential. Note that 15% of our residue pairs falling into the published OCCM structure were long distance (> 30 Å, Supplementary Fig. 6c). This indicates that CLMS unveils dynamic aspects of protein complex topology also in regions of the structure accessible to cryo-EM as will become even more evident in our proteasome analysis.

We next analysed an affinity-purified 26S proteasome sample, containing more than 600 proteins (Supplementary Data 3). The results of our workflow compare favourably with the largest analysis reported on this complex to date^6^ in terms of numbers (n=1644 versus 447 unique residue pairs in the proteasome at 5% FDR) (Fig. 2d,e, Supplementary Fig. 7a, Supplementary Data 1 and 2) and in terms of agreeing with the structure of the individual subunits (6% versus 26% long-distance links (> 30 Å)) (Fig. 2d,e). Links between proteins (n=602) reveal a large amount of topological variability in the proteasome, with 30% (n=179) being not covered by current cryo-EM based models and thus extending our awareness of the proteasome structure to more dynamic regions. Long distance links (n= 191 between and 85 within proteins) are mainly distributed in the base of the proteasome, where ATP-binding and hydrolysis leads to a large conformational variety (Fig. 2f). Indeed, some of these links (n=78) not matching to one structure of the proteasome mapped well to alternative conformational states stabilised by ATP analog^30,31^. State-specific crosslinks were found predominantly in the AAA-ATPase dependent heterohexameric ring (Supplementary Fig. 7) indicating rearrangement of Rpn5 relative to Rpt4 (Fig. 2g). In the s2 state our data support Rpn1 being translated and rotated to be positioned closer to the AAA-ATPase (Fig. 2g). Crosslinks therefore support in solution the cryoEM-based model of substrate transfer to the mouth of the AAA-ATPase heterohexameric ring^30^ and point towards the existence of additional conformational states that remain to be defined to fully understand the complex’s function and that may offer as conformer-specific interactions prime intervention points for small molecule inhibitors.

To probe our 12-fraction protocol in large-scale CLMS we analysed seven high-molecular weight fractions of human cytosol. We identified 3572 unique residue pairs (5% FDR, 528 proteins, Supplementary Fig. 8a, Supplementary Data 1 and 2). This is in line with recent studies reporting 1663 and 3045 unique residue pairs, respectively, albeit using a cleavable crosslinker^4,7^. Interestingly, the overlap between the published data and ours is strikingly low (Fig. 2h) suggesting the approaches are largely complementary.

Our protein-protein interaction network included previously observed complexes like the Mcm2-7 complex, the 26S proteasome, the ribosome, the COPI complex, the TRiC-CCT complex and the HS90B-CDC37-Cdk4 complex (Fig. 2i and Supplementary Fig. 8b-g, 9). For the 26S proteasome we were able to distinguish between different states defining flexibility in the AAA-ATPase ring (Supplementary Fig. 8 g-k)^32^. This indicates the ability of our protocol to unveil dynamic interactions in mixtures nearing the native environment complexity of proteins.

To analyse the mass spectrometric data of these and other studies^4,6,7^ we developed our database search software Xi (Fig 3a, Supplementary Fig. 11, http://xi3.bio.ed.ac.uk/downloads/xiSearch/). The algorithm of Xi has been described conceptually before^33^. It follows an approach that computationally unlinks crosslinked peptides and by doing so circumvents the n2 database problem of crosslinking. Like pLink^9^, StavroX^10^ and Kojak^11^, Xi allows to search any crosslink and protease specificity, thus Xi’s performance was assessed against these three alternatives. StavroX became non-responsive and was not pursued further. Kojak was paired with PeptideProphet^34^ and Xi with xiFDR^35^ to maximise results and control the error rate. We assess the error on the level of unique residue pairs – as this is the actual information of interest. This is native to XiFDR while for Kojak(+PeptideProphet) and pLink the output was sorted by score on the level of PSMs, only the best scoring PSM per residue pair was kept and a 5% FDR on the, now unique residue pairs calculated. By default, XiSearch weights the likelihood of a K versus S, T or Y being involved in a crosslink higher to reduce the number of unique links without strong support by data. For the purpose of comparison XiSearch was run with this feature enabled (marked Xi*) and with this feature disabled, as none of the other tools are supporting a similar consideration. Xi reports 91% more unique links than Kojak+PeptideProphet and 45% more than pLink (Fig. 3b, Supplementary Fig. 10). Note that we and the developers of these tools could not pair Kojak with PeptideProphet for sequential digestion. To check the reliability of residue pairs uniquely reported by Xi, we assessed them for the structurally rigid proteasome core-particle. Less than 3% were found to be long distance and thus very plausibly false, which is in good agreement with the expected FDR of 5%. In summary, Xi performed very favourably compared to other universal software for CLMS.

Our integrated workflow utilises standard crosslinkers, without special chemistries to assist analysis. While recent large-scale studies have successfully used MS-cleavable crosslinkers^7,36^, our work uses standard crosslinkers at no obvious disadvantage. MS-cleavable crosslinkers have yet to be combined with high-density crosslinking and are likely incompatible with crosslinking by non-canonical amino acids^36^ motivating efforts in keeping crosslink chemistry and analysis workflows separate. Our protocol supports this drive and provides a concise, universal protocol to increase data density and ease of use for CLMS in diverse applications that include the detection of dynamic protein interaction regions and topologies which are notoriously difficult to detect using conventional structural biology methods yet are prime therapeutic intervention points.

## Supporting information

XiFDRFileDescription

XiSearchResultFileDescription

## Acknowledgements

We thank Swantje Lenz for the technical support and helpful discussions and Petra Ryl for sample preparation support. We thank Pietro Roversi and CASP12 organizers for supplying UGGT sample and Alberto Riera for contributing OCCM sample. M.L.M. was supported by The International Post-Doc Initiative - IPODI, co-funded by the European Union. This work was supported by the Einstein Foundation, the DFG [RA 2365/4-1], and the Wellcome Trust through a Senior Research Fellowship to J.R. [103139], an Investigator Award to C.S. [107903/Z/15/Z] and a multi-user equipment grant to J.R. [108504], the Biotechnology and Biological Sciences Research Council UK to C.S. [BB/N000323/1] and the Medical Research Council UK to C.S. [MC_U120085811]. The Wellcome Centre for Cell Biology is supported by core funding from the Wellcome Trust [203149].

## Author contribution

M.L.M, L.F. and J.R. developed the study; M.L.M, Z.A.C., F.O’R., A.B., M.B. and M.R.E. performed sample preparation; M.L.M. performed LC-MS analysis; M.L.M., Z.A.C., F.O’R., A.B., T.D., M.B., C.S., S.G. and M.B.S. performed data analysis; L.F. developed Xi; C.W.C. and M.G. provided critical support in data analysis; M.B., C.S., M.R.E. and W.B. contributed with data interpretation; M.L.M., F.O’R., M.B.S., L.F. and J.R. wrote the article with critical input from all authors.

## Materials and Methods

### Reagents

Catalase, Myoglobin, Cytochrome C, Lysozyme, human Serum Albumin (HSA) and Conalbumin were purchased from Sigma. Creatine Kinase was purchased from Roche. C3b was purchased from Complement Technology, Inc. UGGT was kindly provided by Pietro Roversi via the *The Critical Assessment of* protein *Structure Prediction* 12^th^ experiment (CASP12) organizers. Bis[sulsosuccinimidyl]suberate (BS^3^), disuccinimidyl suberate (DSS), sulfosuccinimidyl 4,4’-azipentanoate (sulfo-SDA) and trypsin were purchased from Thermo Scientific. GluC, chymotrypsin and AspN, were purchased from Promega. All media, supplements and phosphate buffered saline (PBS) for cell culture were purchased from PAA Laboratories.

### Sample preparation, crosslinking and digestion with trypsin

The seven standard proteins catalase, myoglobin, cytochrome C, lysozyme, creatine kinase, HSA and conalbumin were resuspended in BS^3^ crosslinking buffer (20 mM HEPES, 20 mM NaCl, 5 mM MgCl_2_, pH 7.8) to a final concentration of 1 mg/ml. Crosslinker was added to a 1:1 (w/w) protein to crosslinker ratio and samples incubated for 2 h on ice. Crosslinking reaction was quenched with excess ammonium bicarbonate (ABC) for 1 h at room temperature (RT). The seven crosslinked proteins were loaded on NuPAGE^TM^ 4-12% Bis-Tris protein gels to isolate the monomeric band of each protein that was then extracted, and in-gel digested with trypsin^38^. After peptide extraction from the gel, peptides from each protein were mixed in a 1:1 weight ratio to a final amount of 200 μg and desalted using C18-StageTips^39^.

For the OCCM complex pUC19-ARS1 beads were used to assemble the OCCM complex as described elsewhere^26^. Crosslinking was performed on beads. BS^3^ was added to 200 μg of the OCCM complex to a 1:8100 protein to crosslinker molar ratio. The sample was incubated for 2 h on ice and the crosslinking reaction was quenched with excess ABC for 1h at RT. The sample was transferred into 8 M urea, reduced with dithiothreitol (DTT), alkylated with iodoacetamide (IAA) and diluted with ABC 50 mM to a final concentration of 2 M urea. Trypsin was added to a protease-to-substrate ratio of 1:50 and the sample was incubated ON at 37°C. Reaction was stopped with 10% (v/v) trifluoroacetic acid (TFA), and the samples divided into four parts and desalted using StageTips.

The 26S proteasome was isolated from *Saccharomyces cerevisiae* by affinity purification using the 3x FLAG-tagged subunit Rpn11 as described elsewhere^40^. For the crosslinking, the 26S proteasome buffer was exchange to BS^3^ crosslinking buffer using 30 kDa molecular weight cut-off (MWCO) filters (Millipore). 200 μg of the 26S proteasome were crosslinked with BS^3^. BS3 was added to a 1:1 (w/w) protein to crosslinker ratio. Samples were incubated for 2 h on ice and the crosslinking reaction was quenched with excess ABC for 1 h at room temperature (RT). The sample was dried using a vacuum concentrator and resuspended in 6 M urea/2 M thiourea for subsequent *in-solution* digestion. Sample was reduced with 2.5 mM DTT for 15 min at 50°C, then alkylated with 5 mM IAA at RT in the dark and diluted with ABC 50 mM to a final concentration of 1 M. Trypsin was added at an enzyme-to-substrate mass ratio of 1:50 and the sample was incubated ON at 37°C. Reaction was stopped with 10% (v/v) TFA, and the samples divided into four parts and desalted using C18-StageTips.

K562 cells (DSMZ, Cat# ACC-10, negatively tested for mycoplasma) were grown in T175 flasks at 37°C in humidified 5% (v/v) CO_2_ incubators in RPMI 1640 media supplemented with 10% (v/v) fetal bovine serum (FBS) + 2 mM glutamine. 3×10^8^ cells were harvested by centrifugation (180xg) and washed 3 times with ice cold PBS. Cells were lysed in lysis buffer (100 mM HEPES pH 7.2, 100 mM KCl, 20 mM NaCl, 3 mM MgCl_2_, 1 mM EDTA, 10% (v/v) glycerol, 1 mM DTT, 10 μg/ml DNAse1, Complete EDTA-free protease inhibitor cocktail, 1 mM PMSF) using a douncer at 4°C. Lysate was cleared of debris by centrifugation at 100,000 x g for 45 min. Native protein complexes were further concentrated by spin filtration using a 100,000 Da cut-off Amicon Ultra centrifugal filter unit (approx. 30 min). For protein co-elution analysis, 100 μl of concentrated lysate (approximately 30 mg/ml) was separated by a Biosep SEC-S4000 (7.8 × 600) size exclusion column on an Åkta Purifier (HPLC) system running at 0.25 ml/min 100 mM HEPES pH 7.2, 100 mM KCl, 20 mM NaCl, 3 mM MgCl_2_, 1 mM EDTA, 10% glycerol. 500 μl fractions were collected. Protein fractions were crosslinked at <1mg/ml (quantified by Bradford) and with 2:1 w/w ratio of BS^3^ to protein. Samples were *in-solution* digested with trypsin as described above.

C3b monomer and dimer were labelled as described elsewhere^17^ and in gel digested with trypsin.

UGGT was crosslinked using Sulfo-SDA using eight different protein to crosslinker ratios (1:0.13, 1:0.19, 1:0.25, 1:0.38, 1:0.5, 1:0.75, 1:1 and 1:1.5 (w/w)). Crosslinking was carried out in two-stages: firstly sulfo-SDA, dissolved in SDA crosslinking buffer (25 μl, 20 mM HEPES-OH, 20 mM NaCl, 5 mM MgCl_2_, pH 7.8), was added to target protein (25 μg, 1 μg/μl) and left to react in the dark for 50 minutes at room temperature. The diazirine group was then photo-activated using UV irradiation, at 365 nm, from a UVP CL-1000 UV Crosslinker (UVP Inc.). Samples were spread onto the inside of Eppendorf tube lids by pipetting (covering the entire surface of the inner lid), placed on ice at a distance of 5 cm from the tubes and irradiated for 20 minutes. The reaction mixtures from the eight conditions corresponding to each experiment were combined and quenched with the addition of saturated ABC (13 μl). Sample was dried in a vacuum concentrator and 200 μg of each protein were *in-solution* digested as described above.

### Parallel Digestion

Parallel digestion with AspN, chymotrypsin and GluC of the seven standard proteins was performed as described for trypsin. After isolating the monomeric bands of each one of the seven standard proteins, those were in-gel digested in parallel with AspN, chymotrypsin and GluC. After peptide extraction from the gel, peptides from each protein digested with the same protease were mixed in a 1:1 weight ratio to a final amount of 200 μg resulting in four samples of the protein standards, one digested with AspN, one digested with chymotrypsin and one digested with AspN. Proteases were added as follows: GluC and chymotrypsin were added to a protease-to-substrate (w/w) ratio of 1:50 and incubated overnight (ON) at 37°C and RT, respectively; AspN was added to a protease-to-substrate ratio (w/w) of 1:100 and incubated ON at 37°C. After parallel digestion samples were fractionated by SEC and analysed by LC-MS/MS.

### Sequential digestion

After trypsin digestion reactions were stopped with 10% (v/v) TFA, divided into four parts and desalted using StageTips. Eluted peptides were dried and while one part, digested with only trypsin, was fractionated by SEC, the remaining three were sequential digested.

Each one of the samples for sequential digestion was resuspended in Ammonium Bicarbonate (ABC) 50 mM and sequential digested as follows: for the sequential digestion with GluC and chymotrypsin, proteases were added to an enzyme-to-substrate ratio of 1:50 and incubated ON at 37°C and RT, respectively. For the sequential digestion with AspN, the enzyme was added at an enzyme-to-substrate ratio of 1:100 and incubated ON at 37°C. After sequential digestion samples were acidified using 10% TFA and the sample volume was reduced to 50 μl by evaporation using a vacuum concentrator. All sequential digested samples were fractionated by SEC.

### Fractionation of peptides by Size Exclusion Chromatography (SEC)

Size exclusion chromatography of crosslinked peptides was performed as described elsewhere^2^. 50 μg of peptides were fractionated in a Shimadzu HPLC system using a Superdex Peptide 3.2/300 (GE Healthcare) at a flow rate of 50 μl/min using SEC buffer (30% (v/v) ACN, 0.1% (v/v) TFA) as mobile phase. Separation was monitored by UV absorption at 215 and 280 nm. Fractions were collected every two minutes over one column volume. The three high-MW fractions were dried, resuspended in 0.1% (v/v) TFA and analysed by LC-MS/MS. All samples in this work, including all the samples used in our proof of principle experiments - replicates, parallel and sequential digestions - of the seven standard proteins, were fractionated by SEC as described in our workflow in Fig 1a.

### In vitro pre-RC assay

The *in vitro* pre-RC assay was performed as described^26,41^. Briefly, ORC (40 nM), Cdc6 (80 nM), Cdt1 (40 nM), MCM2-7 (40 nM) or MCM2-7 ΔC2 (40 nM) were incubated with 6 nM of DNA containing the ARS1 sequence in 50 μl of pre-RC buffer (50 mM HEPES pH 7.5, 100 mM potassium glutamate, 10 mM magnesium acetate, 50 μM zinc acetate, 3 mM ATP, 5 mM DTT, 50 μM EDTA, 0.1% triton-X, 5% glycerol). After 20 minutes at 24 °C, the reactions were washed three times with low salt buffer (pre-RC buffer) or high salt buffer (pre-RC buffer plus 300 mM NaCl) before digestion with 1 U of DNaseI in pre-RC buffer plus CaCl_2_.

### Yeast lethality assay

Yeast strain AS499 (MATa. bar1Δ, leu2-3,-112, ura3-52, his3-Δ200, trp1-Δ-63, ade2-1 lys2-801, pep4) was transformed with pESC-LEU-MCM2-MCM7, pESC-TRP-MCM6-MCM4 and pESC-URA-HA-MCM3-MCM5 (wild type, YC119) or pESC-LEU-MCM2 (Δ848-868)-MCM7 (MCM2-7ΔC2, YC388). Yeast strains YC119 and YC388 were plated on a dropout synthetic complete (SC) medium and incubated at 30 °C for 48 hours. Cells were grown in suspension to 10^7^ cells/ml. Three microliters of a five-fold serial dilution were spotted onto selective plates containing either 2% galactose or glucose. Plates were incubated at 30 °C for 3-5 days.

### LC-MS/MS

Samples were analysed using a Thermo Scientific Dionex Ultimate 3000 RSLCnano system coupled to a Thermo Scientific Orbitrap Fusion Lumos Tribrid mass spectrometer equipped with an EASY-Spray source. Mobile phase A consisted in 0.1% (v/v) FA in water and mobile phase B consisted of 80% (v/v) ACN and 0.1% (v/v) FA in water. Peptides were loaded into a 50 cm EASY-Spray column operated at a temperature of 45°C at a flow rate of 300 nl/min and separated at 300 nl/min using the following gradient: 2% mobile phase B (0-11 min); 2-40% mobile phase B (11-150 min); 40-95% mobile phase B (150-161 min); 95% mobile phase B (161-166 min); 95-2% mobile phase B (166-185 min).

MS data were acquired using a “high-high” acquisition method using the Orbitrap to detect both MS and MS/MS scans. The instrument was operated in a data dependent mode with a cycle time of 3 s. MS1 scans were acquired at a resolution of 120,000 using a scan range from 300 to 1700 m/z and AGC target of 2.5×10^5^ with a maximum injection time of 50 ms. The Monoisotopic Peak Determination (MIPS) was activated and only precursors with charge states between 3 and 8 with an intensity threshold higher than 5.0×10^4^, were selected for fragmentation. Selected precursors were fragmented by HCD using a collision energy setting of 30%. MS2 spectra were acquired at a resolution of 15,000 and AGC of 1.0×104 with a maximum injection time of 35 ms. Dynamic exclusion was set to 60 s after 1 count.

### Data analysis

Thermo raw data were pre-processed using MaxQuant (v 1.5.7.4) to extract the peak list files (APL format). Partial processing in MaxQuant was performed until step 5 with the parameters set to default with exception of the “FTMS top peaks per Da interval” which was set to 100 and no FTMS de-isotoping was allowed. apl files were uploaded to Xi for identification of crosslinked peptides (Xi software is available from http://xi3.bio.ed.ac.uk/downloads/xiSearch/ and the code is available from https://github.com/Rappsilber-Laboratory/XiSearch). For the crosslinking search the parameters used were as follows: MS accuracy, 6 ppm; MS/MS accuracy, 20 ppm; enzyme, trypsin, trypsin+AspN, trypsin+chymotrypsin or trypsin+Gluc depending on the sample digestion conditions; missed cleavages allowed, 4. For the BS^3^ crosslinked samples, carbamidomethylation of cysteines was set as a fixed modification and oxidation of methionine and the crosslinker alone (mass modification: 109.0396 Da), hydrolysed (BS3-OH: 156.0786 Da) or amidated (BS3-NH2: 155.0964 Da) were set as variable modifications. Reaction specificity for BS^3^ was assumed to be with Lysine, Serine, Threonine and Tyrosine or the protein N-terminus. For SDA crosslinked samples, carbamidomethylation of cysteines, oxidation of methionine and the crosslinker alone (mass modification: 109.0396 Da), hydrolysed (SDA-OH: 100.0524 Da) or crosslinker loop (SDA-loop: SDA crosslink within a peptide that is also crosslinked to a separate peptide, 82.0419 Da) were set as variable modifications. Reaction specificity for SDA was assumed to be with Lysine, Serine, Threonine and Tyrosine or the protein N-terminus on one end of the spacer and with all residues on the other end. Apl files were searched against the following databases: for the Standard Protein Mix a database was built containing the sequences corresponding to the crystal structures of the standard proteins used in the mix (PDB accession: 3j7u, 5d5r, 3nbs, 1dpx, 2crk, 1ao6 and 1ova); for C3b we used the fasta corresponding to the Uniprot accession P01024; for the OCCM complex a database containing all the 14 OCCM subunits was used; for the 26S proteasome a linear search of the sample was first performed using a complete *S. cerevisiae* database and the 1% most intense proteins accordingly to the iBAQ value were selected to build the database for the crosslinking search; for each one of the cytosolic SEC fractions the procedure was as for the 26S proteasome, using as database for the linear searched the complete human database; for the UGGT, the uniprot sequence of the protein was used (G0SB58). FDR was estimated on a 5% residue level, including only unique PSMs and boosting results, using xiFDR. Necessary alignments were made using the score matrix BLOSUM62.

Cleavage site protection was calculated by dividing the mean of available miss cleavages for the second protease in the sequential digestion datasets by the mean of the available miss cleavages in the trypsin dataset.

For software comparison, the 26S proteasome was searched using Xi (version 1.6.731), Kojak (version 1.5.5) and pLink (Version 2.3.2) using the same parameters as described before with minor alterations: Searches were performed using 2 missed cleavages, MS accuracy 3 ppm and MS/MS accuracy 20 ppm. For Xi, we used missing mono-isotopic peaks 3. For Xi, Kojak and pLink we gave the same preference for linkage sites in K, S, T, Y.

The mass spectrometry proteomics data have been deposited to the ProteomeXchange Consortium (http://proteomecentral.proteomexchange.org) via the PRIDE partner repository^42^ with the dataset identifier <PXD008550>.

PRIDE reviewer account details: **Username:** **Password:** pNV2lojo

