## Supplementary Information for "An integrated workflow for crosslinking mass spectrometry"


Data are available via ProteomeXchange with identifier PXD008550

### Supplementary figures

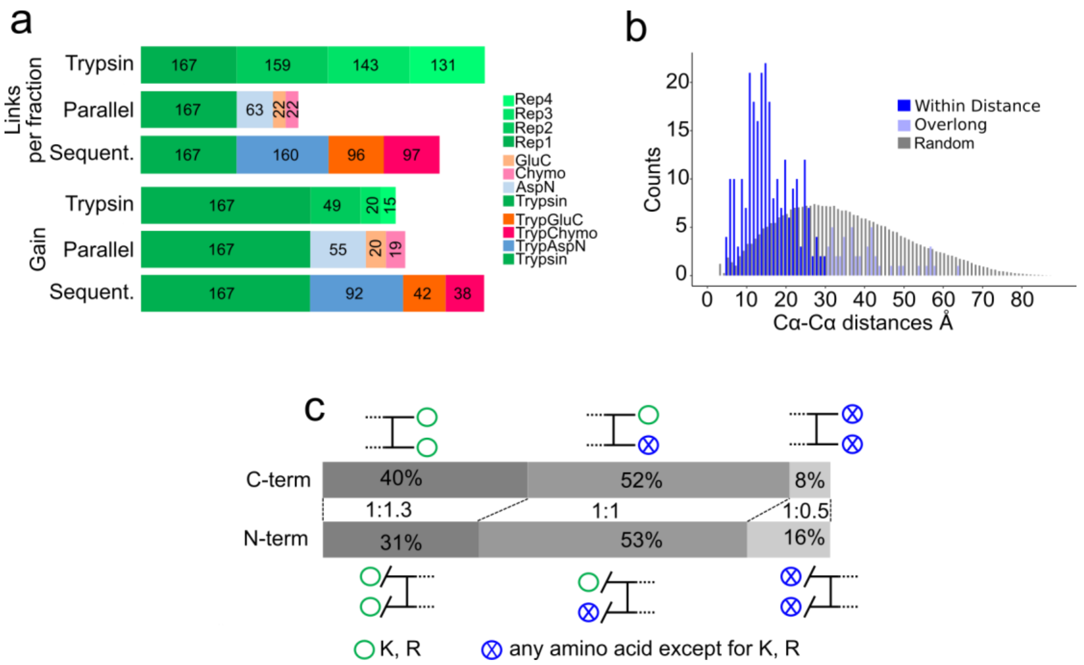


**Supplementary Figure 1:** Sequential digestion increases the number of identified unique residue pairs in a seven-protein mixture. (a) Links per fraction and gain for sequential digestion and the control experiments composed by an experiment using trypsin alone in four replicate and individual digestions with trypsin, AspN, chymotrypsin and GluC. Trypsin yields the higher number of links per sample followed by sequential digestion and individual digestions. However, sequential digestion yields the largest number of unique residue pairs when combining the data. (b) Histogram of the Cα-Cα distances, for 5% FDR. (c) Sequential digested crosslinked peptides show a bias towards having C-termini that end in K or R, which is not the case for the N-termini, showing that sequential digestion delivers smaller tryptic peptides easier to identify by LC-MS/MS.

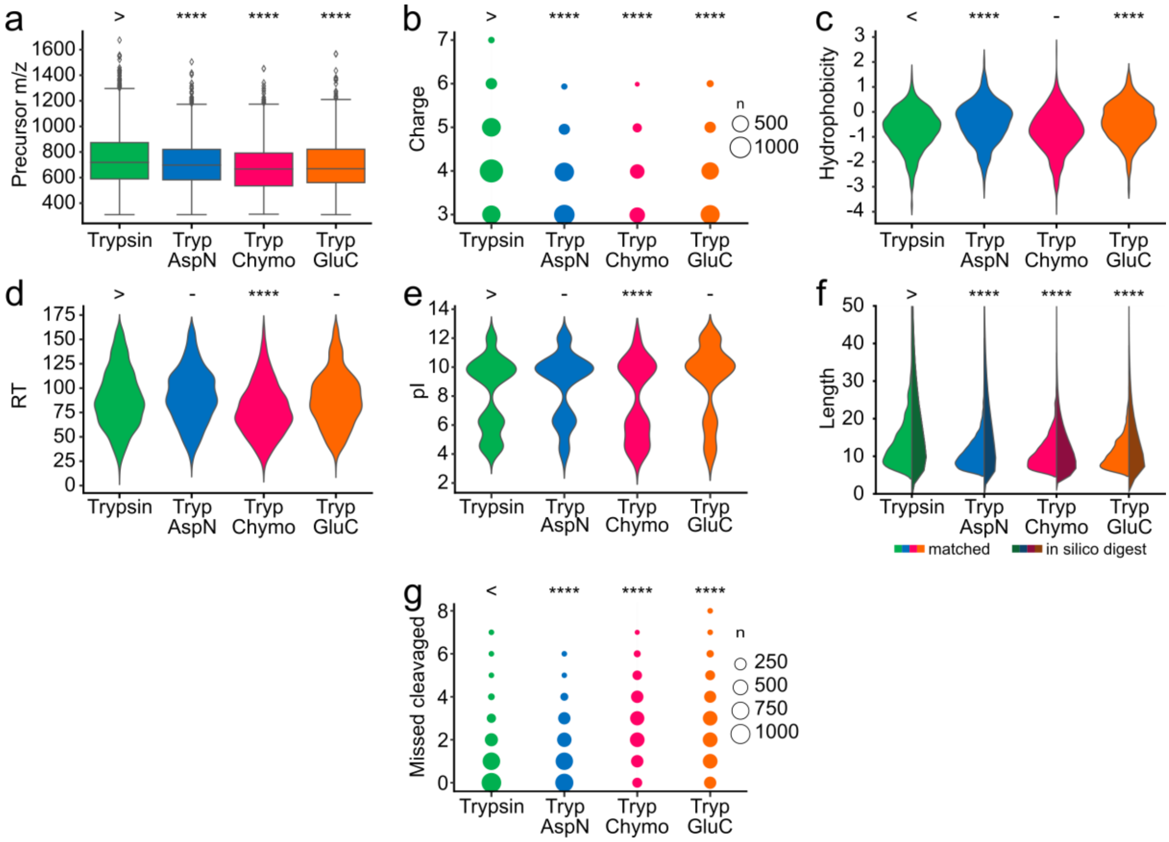


**Supplementary Figure 2:** Properties of crosslinked peptides (i.e. the two linked peptides are considered together) for a seven-protein mixture. For each digestion condition we plotted (a) precursor m/z, (b) observed charge state, (c) calculated hydrophobicity, (d) observed retention time (RT), (e) calculated pI, (f) peptide length, of both, the observed peptides as part of a crosslink (left) and the number of unique crosslinkable peptides resulting from *in-silico* digestion (right), and (g) number of missed cleavages. As expected, sequential digested peptides are smaller and have lower charge states. Sequential digested samples with trypsin+chymotrypsin and trypsin+GluC show more miss cleavages than the other fractions. For statistical testing a one-sided Mann-Whitney-U-test with continuity correction was used. All tests were carried out with trypsin as reference. The sign above the trypsin data (> or <) shows the direction of the alternative hypothesis. (****: 0.0001, ***: 0.001, **: 0.01, *: 0.05, -: 0.1).

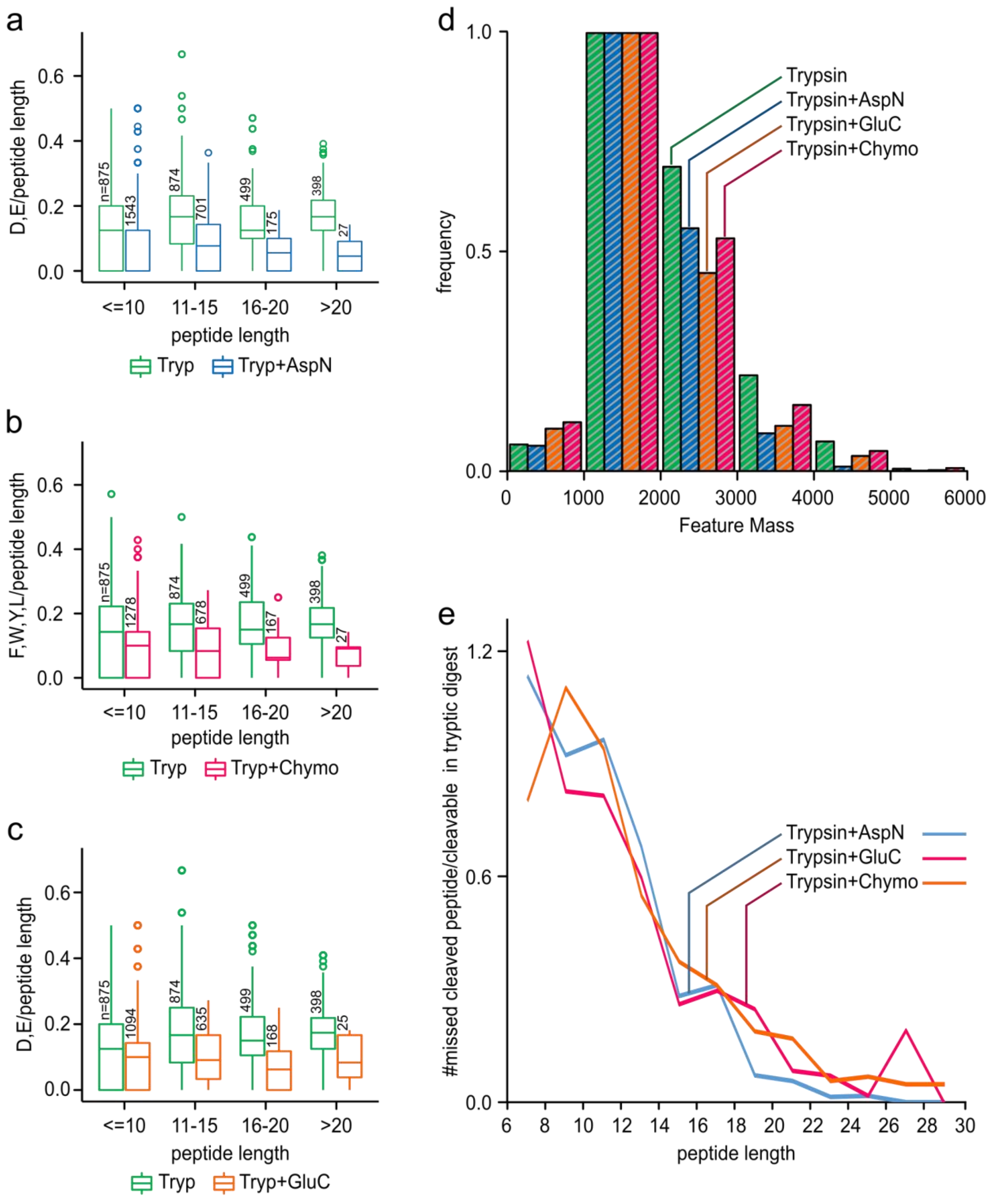


**Supplementary Figure 3:** Cleavage site protection in a BS$^3$-crosslinked 26S proteasome sample. (a, b, c) We determined the number of available cleavage sites for AspN, chymotrypsin and GluC in both the trypsin dataset and their respective datasets (Tryp+AspN, Tryp+Chymo and Tryp+GluC, respectively). Boxplots show that the bigger the observed peptide is, the lower is its number of remaining cleavage sites, showing that large peptides with a higher number of cleavage sites were digested. In turn, smaller peptides contain a larger density of missed cleavage sites thereby indicating that short length protects peptides from digestion. (d) Histogram of intensity weighted MS features as detected by MaxQuant for each digest. The sequential digests show a slight shift to lower masses, but most observed masses are between 1000 and 2000 Da. (e) Protection of peptide from secondary cleavage measured as the number of missed-cleaved peptides in sequential digest divided by the number of peptides in the trypsin digest with potential cleavage sites for the second enzyme.

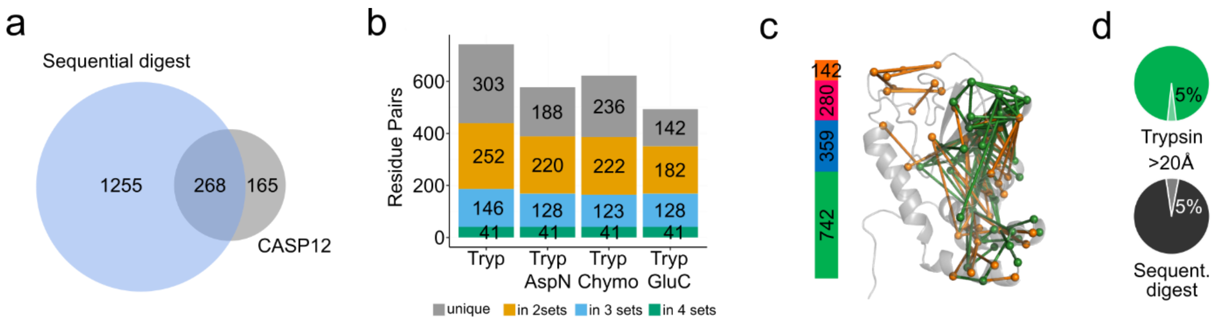


**Supplementary Figure 4:** High-density CLMS applied to CASP12 target UGGT (PDB|5NV4), using sequential digestion and Xi. (a) Comparison between sequential digestion and CASP12 results. Sequential digestion provides a higher number of unique residue pairs. (b) Overlap of the number of residue pairs for the four digestion conditions in the sequential digestion strategy. Sequential digestion complements trypsin digestion increasing the number of identified unique residue pairs. (c) Sequential digestion workflow applied to high-density CLMS shows a gain by a factor of almost two by adding AspN (blue), chymotrypsin (pink) and GluC (orange) to trypsin (green). Additional residue pairs identified upon sequential digestion (in orange) cover regions of the protein structure not observed with trypsin (in green). (d) The percentage of long distance residue pairs (> 20 Å) is 5% for both trypsin and sequential digestion, in agreement with our calculated FDR.

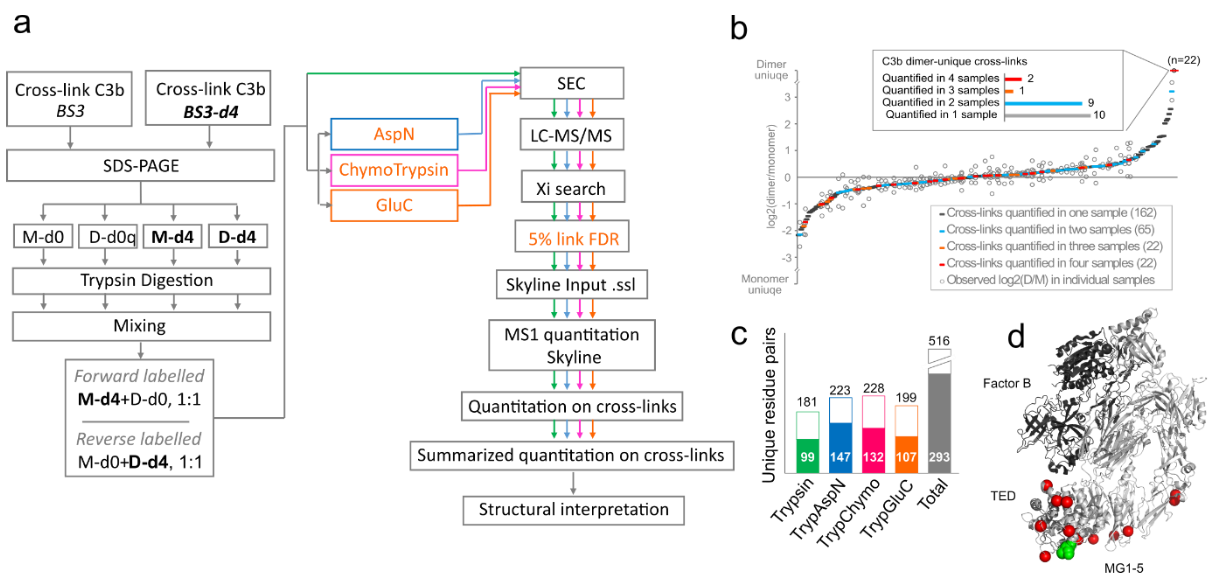


**Supplementary Figure 5:** Sequential digestion applied to QCLMS of C3b dimerization (a) QCLMS-sequential digestion strategy. A C3b sample was divided into two aliquots and crosslinked with either bis[sulfosuccinimidyl] suberate ($BS^3$) or its deuterated analogue bis[sulfosuccinimidyl] 2,2,7,7-suberate-d4 ($BS^3$-d4). BS3 and BS3-d4 crosslinked C3b samples were then separately subjected to SDS-PAGE and the monomer and dimer bands of BS3 and BS3-d4 crosslinked C3b were excised and in-gel digested. Based on C3b abundance, BS3 crosslinked C3b monomer and BS3-d4 crosslinker dimer samples were 1:1 mixed (named as forward-labelled). BS3-d4 crosslinked C3b monomer and BS3 crosslinker dimer samples were also mixed in 1:1 ratio as a reserve-labelled replica. Each of them was divided into four aliquots from which three were further digested each with a second protease. All digested samples were fractioned by SEC chromatography and analysed by LC-MS/MS. Searches were performed by Xi and confidence of identified residue pairs assessed by xiFDR. Quantitation was performed using Skyline. (b) Quantitation reproducibility. The use of different proteases did not lead to major changes on quantified dimer to monomer signal ratios. (c) Half of the identified residue pairs per fraction were quantified. (d) C3b (light grey) bound to factor B (dark grey) (PDB|2XWJ) with position of thioester (green) and position of C3b dimer-specific links (red).

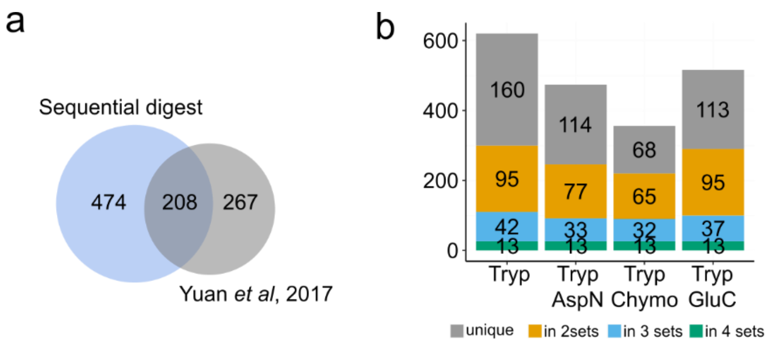


**Supplementary Figure 6:** The OCCM complex analysed by sequential digestion and Xi (a) Comparison between sequential digestion workflow and literature results. Sequential digestion provides a higher number of unique residue pairs. (b) Overlap of the number of residue pairs for the four digestion conditions in the sequential digestion strategy. Sequential digestion complements trypsin digestion increasing the number of identified unique residue pairs. (c) Gain of unique residue pairs by using sequential digestion. The bar represents the number of unique residue pairs gained by adding AspN (blue), chymotrypsin (pink) and GluC (orange) to the digestion with trypsin (green). (d) The percentage of long-distance residue pairs (> 30 Å) is the same for both trypsin and sequential digested samples (18%). The relatively large proportion of long-distance links is explained by the flexible structure of the OCCM complex.

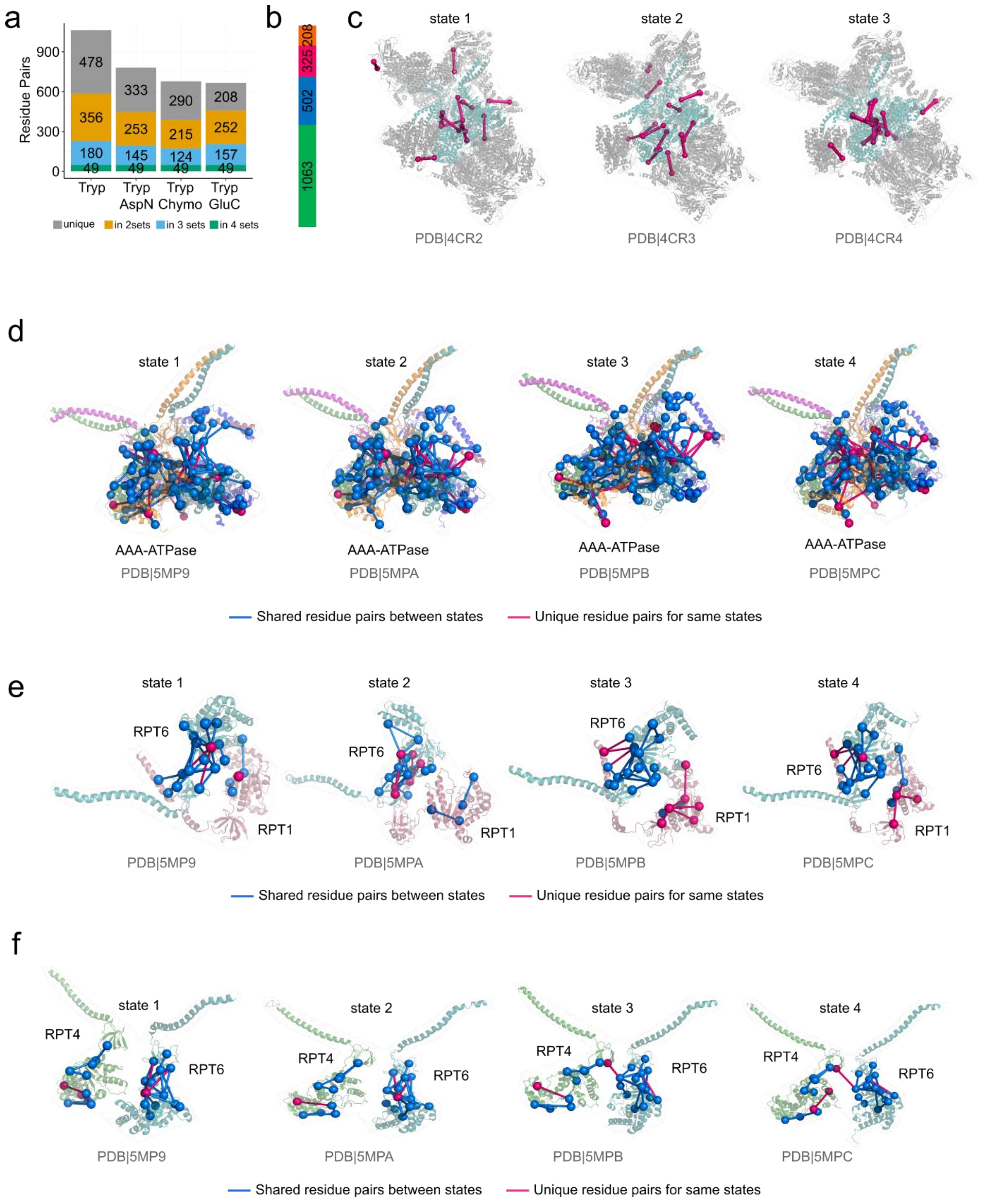


**Supplementary Figure 7:** The different states of the 26S Proteasome. (a) Overlap of the number of residue pairs for the four digestion conditions in the sequential digestion strategy. Sequential digestion complements trypsin digestion increasing the number of identified unique residue pairs. (b) Gain of unique residue pairs by using sequential digestion. The bar represents the number of unique residue pairs gained by adding AspN (blue), chymotrypsin (pink) and GluC (orange) to the digestion with trypsin (green). (c) Links matching the different states of the 26S proteasome described by Unverdorben *et al*. were mapped to the respective structures. (d) Links matching the different states of the AAA-ATPase dependent heterohexameric ring from the 26S proteasome described by Wehmer et al, 2017 were also mapped to the respective structures. (e, f) A closer look to the proteins RPT6 and RPT1 and RPT4 and RPT6, shows the structural rearrangements of the AAA-ATPase dependent hererohexameric ring throughout the four states.

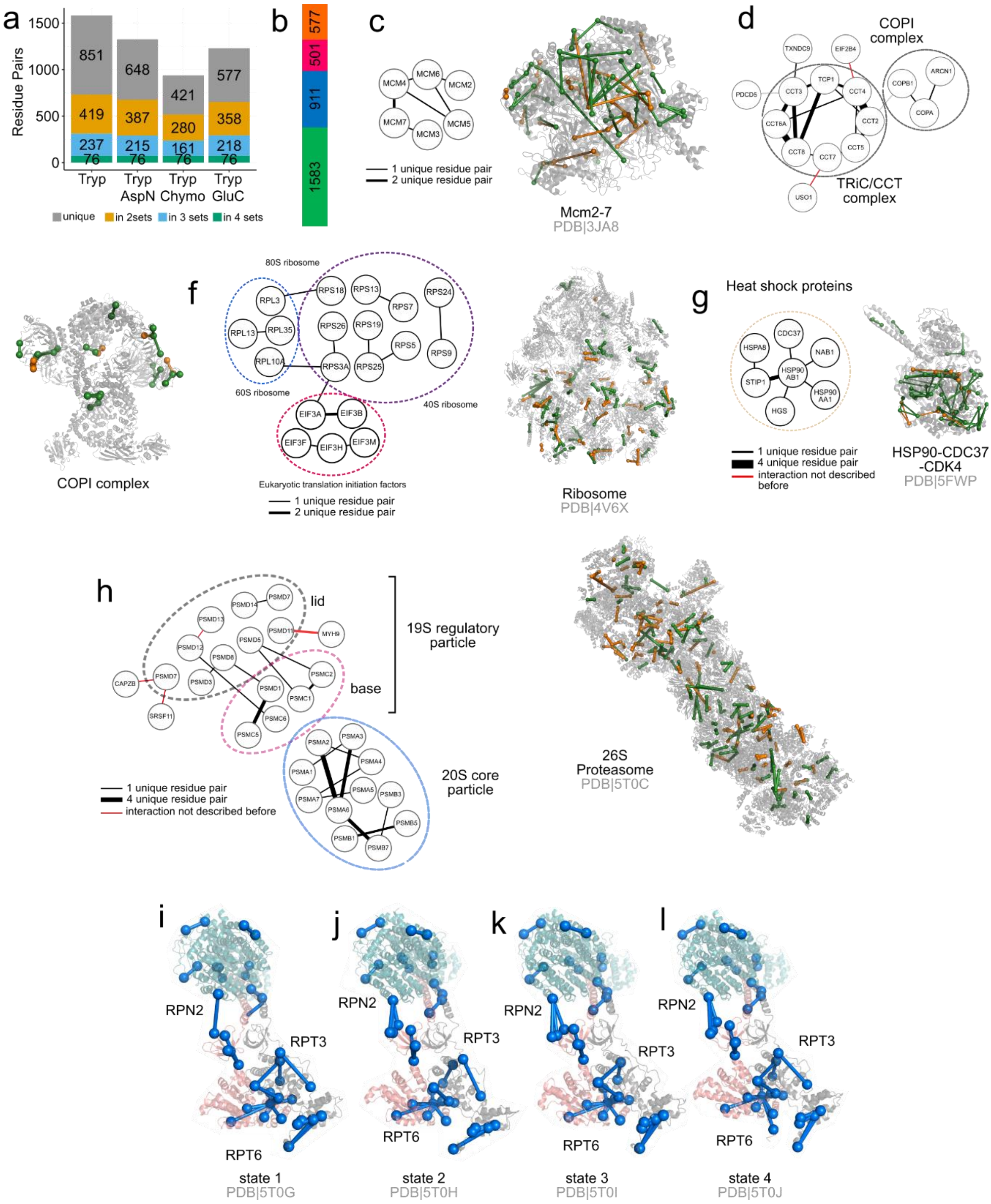


**Supplementary Figure 8:** CLMS of a human cytosol as a complex mixture using sequential digestion and Xi. (a) Overlap of the residue pairs obtained by the four digestion conditions applied. Sequential digestion complements trypsin digestion increasing the identified unique residue pairs. (b) Gain of unique residue pairs by using sequential digestion. The bar represents the number of unique residue pairs gained by adding AspN (blue), chymotrypsin (pink) and GluC (orange) to the digestion with trypsin (green). (c, d, e, f, g, h) Among others, we identified unique residue pairs for the MCM2-7 complex, the TriC/CCT complex the COPI complex, the ribosome, the HSP90-CDC37-CDK4 complex and the 26S proteasome. The respective protein-protein interaction network is displayed and identified residue pairs were mapped into the respective crystal structures. Tryptic links are displayed in green and non-tryptic links are displayed in orange. (i, j, k, l) Despite the complexity of the sample we were able to identify the four states of the 26S proteasome showing the flexibility of the AAA-ATPase dependent heterohexameric ring.

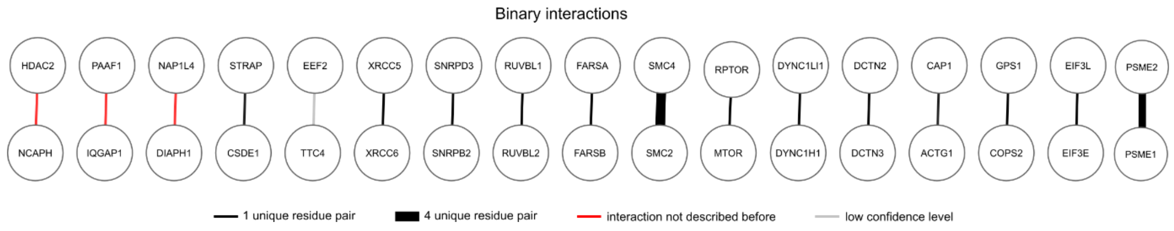


**Supplementary Figure 9:** Binary interactions for the cystosol of human cells. Apart from the identified networks, several binary interactions were also found in the cytosol of human cells represented in this figure.

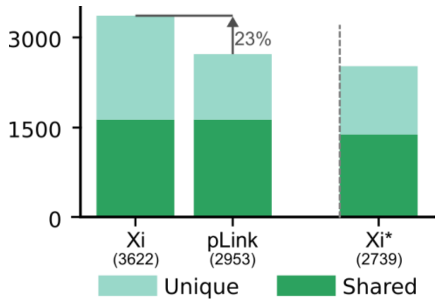


**Supplementary Figure 10:** Comparison of the sequential digest protocol when combined with XiSearch or pLink. Search was done with BS3 specificity defined to be Lysine, Serine, Threonine and Tyrosine. Xi searches where run twice: once treating all linkable residues equal (as is done in pLink) and once (denoted Xi*) with giving priority to Lysine over Serine, Threonine and Tyrosine. A consequence of Xi* is that reporting a crosslink involving one of the side-reactions in favour of a near-by Lysine requires a substantial amount of evidence.

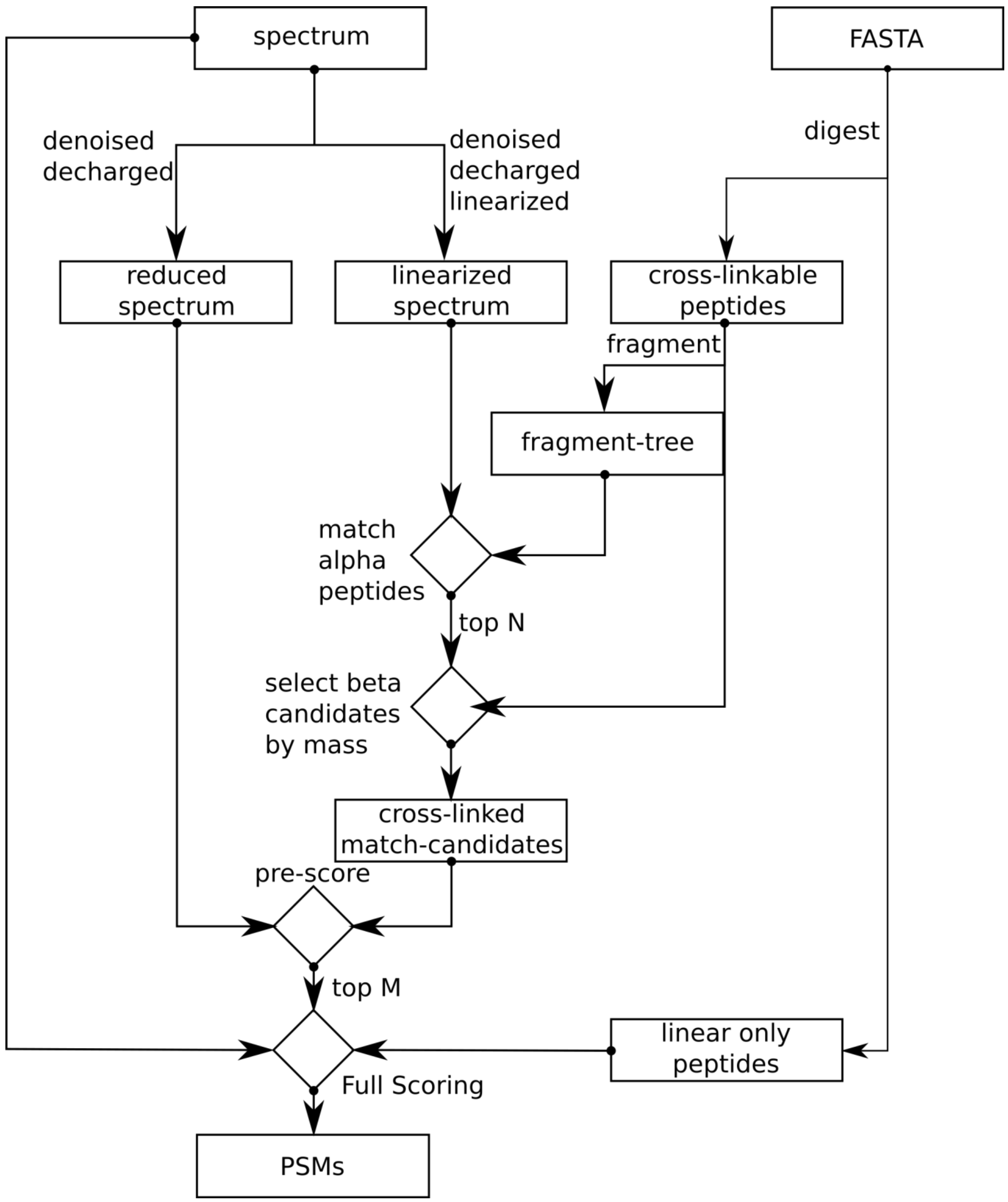


**Supplementary Figure 11:** XiSearch follows a three-step approach. In the first step it tries to identify peptides with an unknown modification that explain the spectrum best. High resolution data helps in this step in two ways. Knowing the charge, and therefore the mass of a fragment, enables to predict whether a fragment is linear or crosslinked[33]. This is important as only linear fragments are used to select peptide candidates. Secondly, with the knowledge that a fragment is probably crosslinked we can invert the crosslinked fragments into their linear counterparts. By doing these two steps we can de-convolute a crosslinked spectrum into a spectrum containing almost exclusively linear fragments of two “independent” peptides. Additionally, the spectrum also gets de-charged and de-noised (linearized spectrum). To enable a fast identification, we create an in-memory representation of all primary fragment(e.g. b- and y-ions)-to-peptide relationships that can be derived from the search-database (fragment-tree). This fragment-tree is then used to identify and quickly score alpha-peptide-candidates in the linearized spectrum while ignoring the precursor mass. Secondly, the top-n candidates are then forwarded to the beta-peptide selection. Here we take for each alpha peptide all peptides that fit the mass-gap between the alpha-peptide candidate plus crosslinker and the precursor mass. The whole list of peptide-pair candidates is then again preliminarily scored. Third and finally the top-m candidate pairs are then fully scored and reported.

### Xi search: step by step instructions

Pre-requirements:

- Ensure java version 8 (64 bit) is installed on your computer (https://www.java.com/ note that when downloading with a 32-bit browser an explicit selection of a 64-bit download is required).
- For large scale analysis a computer with 16 GB memory is recommended.
- Operating system: Windows (Windows Server 2008 and 2012 and Windows 8 and 10) and Linux (Debian 9, Ubuntu 14.04, Ubuntu 16.04) were tested successfully – but expect that any system running java 8 (64 bit) should suffice.

Installation:

1) Download the ZIP file from http://xi3.bio.ed.ac.uk/downloads/xiSearch/.
2) Extract the compressed files to your computer (e.g. C:\Users\John). You will have a new folder (C:\Users\John\XiSearch).
3) If you are using Windows start the *startXiWindows* file. If you are using Unix start the *startXiUnix* file. This will start Xi with a simple graphical user interface (GUI), and the actual java library (XiSearch.jar).
4) The GUI has 5 tabs. In the first tab *Run*, choose the location where you want to save your results under the box *result*. Optional: In *peak annotations* you can define a file for writing out full annotations of the spectra.

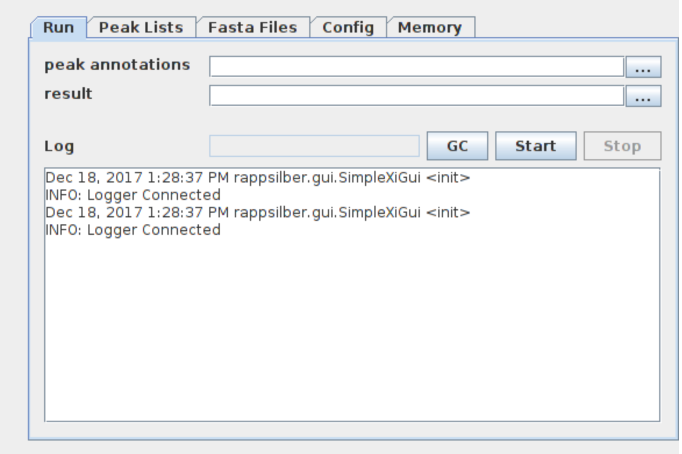

5) In the second tab *Peak Lists* select the peak lists to be searched (Supported formats: mgf-files as generated from msconvert or apl-files as produced by MaxQuant. For mgf-files of other sources the config may need to be adapted to define how to read out a run-name and a scan-number from the TITLE= tags in the files).

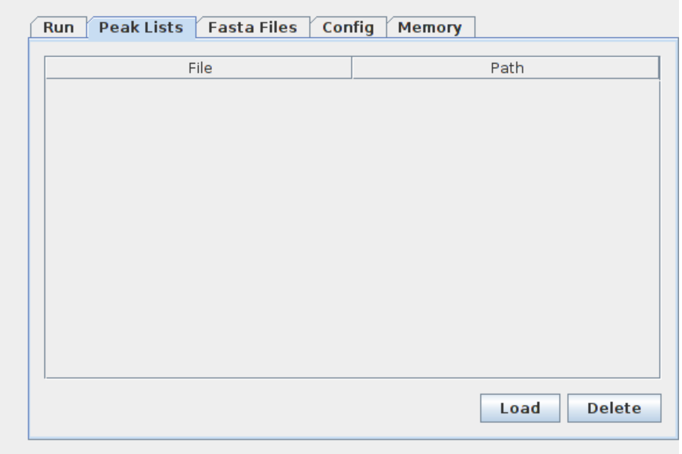


6) In the third tab *Fasta Files* select the FASTA files to be searched. *Load* opens a file selection dialog. Multiple files can be selected there from the computer and one can be defined as custom decoy database by checking the appropriate box. If no decoy database is given, the program automatically generates one based on the target database.

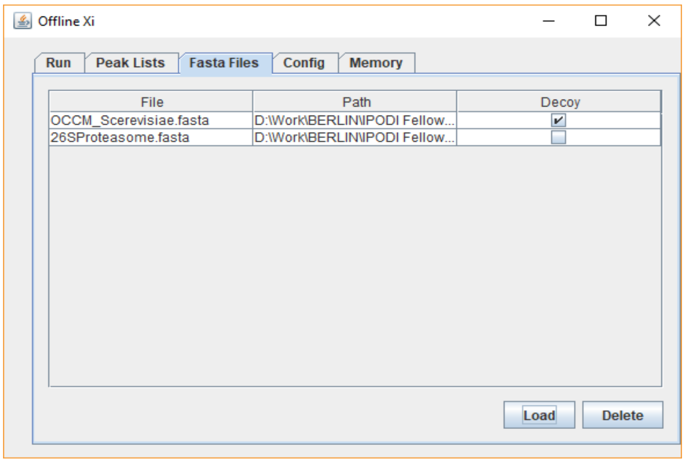


7) In the fourth tab *Config* the search is configured. All parameters are defined in form of a text file that can be edited in the GUI. The tab opens displaying a default config as an example that defines a search with BS3 as crosslinker, carbamidomethylation on cysteines as fixed modification, oxidation of methionine as variable modification and trypsin as the protease of choice. Additional options are still included with a short description of their function, example and the syntax. When you have your config file defined you can save it for further searches under the box *Save*. *Load* allows you to open saved config files from your computer.

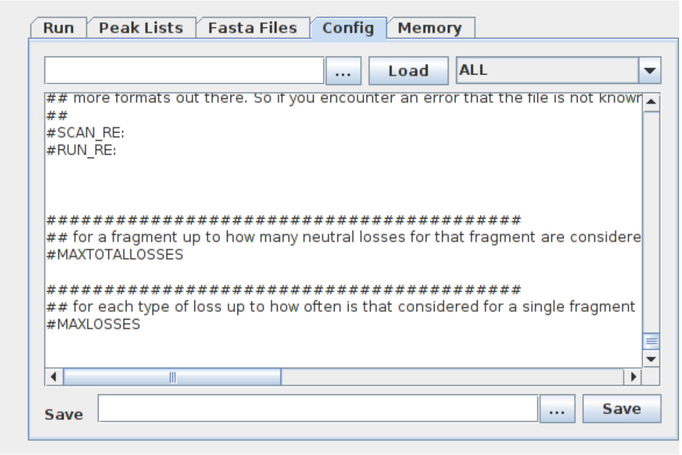


8) Go back to the first tab *Run* and press the *Start* button to start your search.

Notes:

- Depending on the size of the sequence-database and the number of search threads the start-script might need to be adapted to permit Xi to use a larger amount of memory (-Xmx option). This should not exceed the amount of free memory available without running Xi. For example, if a computer has 8 GB of RAM but per default 5 are used by other programs then Xi should get at most 3 GB as otherwise part of the program is likely to be swapped out to disk and in effect will run extremely slow.
- Additionally to the GUI a command line interface is provided that can be used via "java –Xmx4g -cp Xlink.jar rappsilber.applications.Xi --config=[config-file] --peaks=[path to peaklist] --fasta=[path to fasta file] --conf=[some xi-parameters] --output=[csv-file] --peaksout=[csv-file] --exampleconfig=[path]" Description of the arguments can be gained by running java -cp Xlink.jar rappsilber.applications.Xi --help
- XiSearch does not assess the FDR. This is done by a separate application XiFDR [35]. XiSearch result files can be directly read into XiFDR.
- An example dataset is supplied together with the Xi download for testing. The folder contains:
    1. An apl file resulting from a $BS^3$-crosslinked sample digested with trypsin.
    2. The corresponding database
    3. A config file (example.conf)
    4. The result file as obtained by XiSearch (ready for input into XiFDR)

Search time for the example dataset using an Intel-Core i7-5500U with 8 GB memory and Windows10 (laptop): approx. 3 hrs.
