## Supplementary material for "An integrated workflow for crosslinking mass spectrometry": XiFDRFileDescription

### XiFDR File Description

#### Summary

Provides a summary on how the fdr estimation was performed.
It has several section:

#### Settings section:

Provides an overview of what settings where used to produce the result

| | | | |
|---|---|---|---|
| xiFDR Version: | **Version of XiFDR that produced the file** | | |
| Source: | **Where the data where read from** | | |
| | Target FDRs: | Minimum supporting peptides | Directional |
| psm | **FDR applied at PSMs** | | false |
| peptide pair | **FDR applied at peptide pairs** | | false |
| protein group | **FDR applied at protein groups** | 1 | |
| Link | **FDR applied at residue pairs** | 1 | false |
| Protein Group Pair | **FDR applied at protein group pairs** | 1 | false |
| max next level fdr factor (report-factor): | deprecated | | |
| minimum peptide length | **Minimum length of peptide to be considered for FDR** | | |
| unique PSMs | | | |
| Accepted ambiguity: | do to ambiguity of peptide source a peptide pair can represent several links | | |
| Links for one peptide pair | **peptide pairs representing more unique links then this will be ignored** | | |
| Protein pairs for one peptide pair | **peptide pairs representing more unique protein pairs then this will be ignored** | | |
| Length-Group: | **PSMs and peptide pairs get split by the length of the shortest peptide** | | |

#### Result Summary:

presents the number of unique matches that passed all FDRs. This including being filtered back to what passed higher level FDRs. Meaning if a PSM-level FDR of 5% was defined and also a peptide FDR of 5% then the PSM-numbers shown here are the ones that first passed the 5%PSM FDR and supported peptide pairs that passed the 5% Peptide Pair FDR. This is given ones for all and split into internal and between links and there into target-target, target-decoy and decoy matches.

| class | all | Internal TT | Internal TD | Internal DD | Between TT | Between TD | Between DD | Linear T | Linear D |
|---|---|---|---|---|---|---|---|---|---|
| Input PSMs | **all psms** | | | | | | | | |
| fdr PSMs | **PSMS passed all FDRs** | **x** | **x** | **x** | **x** | **x** | **x** | **x** | **x** |
| fdr Peptide Pairs | **peptide pairs passed all FDRs** | **x** | **x** | **x** | **x** | **x** | **x** | **x** | **x** |
| fdr Link | **residue pairs passed all FDRs** | **x** | **x** | **x** | **x** | **x** | **x** | | |
| fdr Protein Group Pairs | **protein pairs passed all FDRs** | **x** | **x** | **x** | **x** | **x** | **x** | | |
| fdr Protein Groups | **protein groups passed all FDRs** | | | | | | | **x** | **x** |

#### Level Information

Then comes one section for each level of information:

It provides the information about what went into the calculation (e.g. what passed the lower level FDRs and prefilter)

| **Level** detailed summary | | | |
|---|---|---|---|
| Group | **Group 1** | **Group 2** | **...** |
| Input | **How many matches where considered in each group** | | |
| TT | **How many target target entries where in the input** | | |
| TD | **How many target decoy entries where in the input** | | |
| DD | **How many decoy decoy entries where in the input** | | |
| passing fdr (0.05) | **How many matches passed the FDR** | | |
| TT | **How many target target matches passed the FDR** | | |
| TD | **How many target decoy matches passed the FDR** | | |
| DD | **How many decoy decoy matches passed the FDR** | | |
| last fdr < 0.05 | **What was the last fdr that did not exceed the target** | | |

| | |
|---|---|
| | **FDR** |
| higher fdr (> 0.05) | **What would have been the next detectable FDR larger then the target FDR** |
| lower fdr (<= 0.05) | **What was the next detectable lower FDR below the “last FDR”** |
| final | **How many matches passed this FDR and higher level FDRs** |

### PSM file

There are two PSM-files generated – one for linear PSMs (non-cross-linked) and one for cross-linked PSMs. Both share the same format but in the linear file all fields pertaining to peptide2 are empty.

Columns:

| | |
|---|---|
| PSMID | **A unique id assigned to a PSM either provided in the input or made up from scan run and peptide sequences** |
| run | **Run Name as provided in the input** |
| exp charge | **experimental spectrum information as provided from the input** |
| exp m/z | |
| exp mass | |
| exp fractionalmass | **the part of the “exp mass” that is behind the dot e.g. exp mass = 734.987 the fractional mass would be 0.987** |
| match charge | **the charge state matched** |
| match mass | **that mass of the peptides + cross-linker** |
| match fractionalmass | **as above for the match-mass** |
| scan | **Scan-number** |
| Protein1 | **accession of protein1 (as in input)** |
| Description1 | **description of protein 1 (as in input)** |
| Decoy1 | **is the first protein a decoy protein** |
| Protein2 | **accession of protein2 (as in input)** |
| Description2 | **description of protein 2 (as in input)** |
| Decoy2 | **is the second protein a decoy protein** |

| | |
|---|---|
| PepSeq1 | **sequence of peptide1** |
| PepSeq2 | **sequence of peptide 2** |
| PepPos1 | **position of peptide 1 in protein1** |
| PepPos2 | **position of peptide 2 in protein2** |
| PeptideLength1 | **length of peptide1 in amino-acids** |
| PeptideLength2 | **length of peptide2 in amino-acids** |
| LinkPos1 | **Link-site in peptide1** |
| LinkPos2 | **link-site in peptide 2** |
| ProteinLinkPos1 | **link-site in protein 1** |
| ProteinLinkPos2 | **link-site in protein 2** |
| Charge | **matched charge** |
| Crosslinker | **the name of the cross-linekr involvet with this match** |
| Score | **score of the PSM** |
| isDecoy | **is this a match involving a decoy peptide** |
| isTT | **is this a target target match** |
| isTD | **is this a target decoy match** |
| isDD | **is this a decoy decoy match** |
| fdrGroup | **under what group was this match considered** |
| lowerFDR | **next lower detectable FDR** |
| fdr | **FDR associated with the match score** |
| higherFDR | **next higher FDR** |

### Peptide Pair

All cross-linked peptide pairs passing all FDRs

| | |
|---|---|
| PeptidePairID | **a numeric ID for the peptide pair** |
| PSMIDs | **all psms that support this peptide pair** |
| Protein1 | **accession of protein1 (as in input)** |
| Description1 | **description of protein 1 (as in input)** |

| | |
|---|---|
| Decoy1 | **is the first protein a decoy protein** |
| Protein2 | **accession of protein2 (as in input)** |
| Description2 | **description of protein 2 (as in input)** |
| Decoy2 | **is the second protein a decoy protein** |
| Peptide1 | **sequence of peptide1** |
| Peptide2 | **sequence of peptide 2** |
| FromSite | **Link-site in peptide1** |
| ToSite | **link-site in peptide 2** |
| FromProteinSite | **link-site in protein 1** |
| ToProteinSite | **link-site in protein 2** |
| psmID | **Legacy – to be deleted** |
| Crosslinker | **the cross-linker for this peptide pair** |
| Score | **the score as derived from PSMs** |
| isDecoy | **is this a match involving a decoy peptide** |
| isTT | **is this a target target match** |
| isTD | **is this a target decoy match** |
| isDD | **is this a decoy decoy match** |
| fdrGroup | **under what group was this match considered** |
| fdr | **FDR associated with the match score** |

### Protein Group

Lists all protein groups that where identified in the sample

| | |
|---|---|
| ProteinGroupID | **a unique id assigned to the protein group** |
| ProteinGroup | **accession numbers of all proteins in this group** |
| Descriptions | **descriptions of all proteins in the group** |
| Sequence | **Amino acid sequences of all proteins in the group** |

| | |
|---|---|
| Crosslinker | **all cross-linker seen involved with the protein group** |
| Score | **a score for the protein group based on the peptide pairs supporting the group** |
| isDecoy | **is this a match involving a decoy peptide** |
| isTT | **is this a target target match** |
| isTD | **is this a target decoy match** |
| isDD | **is this a decoy decoy match** |
| PSM IDs | **list of all PSM-IDs supporting the Protein group** |
| fdrGroup | **under what group was this match considered** |
| fdr | **FDR associated with the match score** |
| File Name1 | **Highest score of a psm supporting this protein group in the given run** |
| File name 2 | |
| ... | |

### Link

All residue pairs that passed all FDRs

| | |
|---|---|
| LinkID | **A unique ID assignt to each link** |
| PeptidePairIDs | **list of supporting peptide pair Ids** |
| PSMIDs | **list of supporting PSM Ids** |
| Protein1 | **Protein 1 that is linked** |
| Description1 | **description of protein1** |
| Decoy1 | **is protein 1 a decoy?** |
| Protein2 | **Protein 2 that is linked** |
| Description2 | **description of protein2** |
| Decoy2 | **is protein 2 a decoy?** |
| fromSite | **link site within protein 1** |
| ToSite | **link site within protein 2** |
| Croslinkers | **cross-linkers that this link was observed with this link** |

| | |
|---|---|
| Score | **score for the link based on all supporting peptide pairs** |
| isDecoy | **is this a match involving a decoy peptide** |
| isTT | **is this a target target match** |
| isTD | **is this a target decoy match** |
| isDD | **is this a decoy decoy match** |
| count PSMs | **how many PSMs support this link** |
| count peptide pairs | **how many unique peptide pairs support this link** |
| fdrGroup | **under what group was this match considered** |
| fdr | **FDR associated with the match score** |
| File Name1<br>File name 2<br>… | **Highest score of a psm supporting this protein group in the given run** |

PPI

List of all protein pairs that passed all FDRs

| | |
|---|---|
| ProteinGroupPairID | **A unique ID assigned to each protein pair** |
| LinkIDs | **list of supporting residue pair Ids** |
| PeptidePairIDs | **list of supporting peptide pair Ids** |
| PSMIDs | **list of supporting PSM Ids** |
| Protein1 | **Protein 1 that is linked** |
| Descriptions1 | **description of protein 1** |
| isDecoy1 | **is protein 1 a decoy?** |
| Protein2 | **Protein 2 that is linked** |
| Description2 | **description of protein 2** |
| isDecoy2 | **is protein 2 a decoy?** |
| Crosslinker | **cross-linkers that this link was observed linking these proteins** |
| Score | **score for the link based on all supporting residue pairs** |
| isDecoy | **is this a match involving a decoy peptide** |
| isTT | **is this a target target match** |
| isTD | **is this a target decoy match** |
| isDD | **is this a decoy decoy match** |

| | |
|---|---|
| count PSMs | **how many PSMs support this link** |
| count peptide pairs | **how many unique peptide pairs support this link** |
| count links | **how many unique residue pairs support this link** |
| fdrGroup | **under what group was this match considered** |
| fdr | **FDR associated with the match score** |
