## Supplementary material for "An integrated workflow for crosslinking mass spectrometry": XiSearchResultFileDescription

### XiFDR Search Results File Description

The XiSearch output has several columns.
The columns fall into several categories:

- general match information
- per peptide information
- sub-scores
- final score

Following the columns are explained in these groups

#### General Match Information

Basic information pertaining the whole PSM.

| *Field* | *Description* |
|---|---|
| Run | Name of the raw-file as passed out from the peak-list |
| Scan | scan number of the spectrum as passed out from the peak-list |
| ScanInputIndex | index of the spectrum in the peak-list |
| Source | name of the actual file containing the spectrum |
| ElutionStart | Start of the elution as passed out from the peak-list |
| ElutionEnd | end of the elution as passed out from the peak-list |
| PrecursorMass | The mass of the precursor as given by the peak-list |
| PrecoursorCharge | The charge of the precursor as given by the peak-list |
| PrecurserMZ | The m/z value of the precursor as given by the peak-list |
| CalcMass | the theoretical mass according to the matched peptides |
| CalcMZ | the theoretical m/z according to the matched peptides |
| validated | did it pass auto-validation (mainly the autovalidation score with some heuristics against some corner cases) |
| decoy | is this a decoy match |
| MatchRank | the rank of the match according to score (mostly) |
| Crosslinker | the name of the cross-linker (empty for linear matches) |
| CrosslinkerMass | the mass of the cross-linker |
| decoyCrosslinker | was the cross-linker found here defined as a decoy cross-linker |

#### Per Peptide Information

Information about each peptide in the PSM. X is the number of the peptide.

| *Field* | *Description* |
|---|---|
| ProteinX | accession number of the protein |
| FastaX | FASTA-header of the protein |
| ProteinXdecoy | is this protein a decoy protein |
| PeptideX | sequence of the peptide |
| BasePeptideX | sequence without any modification |
| PeptideLinkMapX | sequence of the peptide with associated linkage score (displaying a likelihood for a residue to be the linkage site) |
| PeptideMassX | Neutral Mass of the Peptide |
| StartX | start of the peptide in the protein |
| LengthPeptideX | length of the peptide |
| LinkX | linkage site in the peptide |
| Linked AminoAcid X | what amino-acid was linked |
| LinkWindowX | a +- 20 amino acid window around the detected protein linkage site |
| ProteinLinkX | where in the protein is the link |
| ProteinCountX | if the peptide was ambiguous in how many proteins was it found in total |
| PositionCountX | if the peptide was ambiguous in how often was it found in total |
| ModificationsX | list of modifications on this peptide |
| ModificationPositionsX | where in the peptides these modification are |
| ModificationMassesX | delta masses of the modifications |
| OpenModPositionX | Legacy column |
| OpenMassX | Legacy column |
| OpenModWindowX | Legacy column |

#### Sub-Scores

XiSearch calculates a range of sub scores. Following is a description of all columns representing these sub-scores.

##### Candidate Sub-Scores

XiSearch does work in a multistage approach – first it generates a list of alpha candidates and from these a list of candidates peptide pairs that are then fully scored. The following scores pertain are derived from the candidate lists.

| *Field* | *Description* |
|---|---|
| betaCount | for the given alpha peptide - how many beta peptides fit by mass |
| betaCountInverse | 1/betaCount |
| mgcRank | rank of the alpha peptide among the list of alpha peptide candidates |
| mgxRank | rank of the peptide pair among the peptide pair candidates |
| mgcScore | candidate score of the alpha peptide |
| mgcDelta | difference to the second best alpha peptide candidate |
| mgcShiftedDelta | deprecated |
| mgcAlpha | =mgcScore |
| mgcBeta | the alpha candidate score for the beta peptide |
| mgxScore | the candidate score for the peptide pair |
| mgxDelta | difference to the second best peptide pair candidate score |
| delta | difference of the full score to the second best peptide pair (disregarding modifications) |
| deltaMod | difference of the full score to the second best peptide pair (considering modifications) |
| combinedDelta | average of delta and main score |
| match score | main score of the PSM (last entry per line) |

#### Mass-Errors

| *Field* | *Description* |
|---|---|
| Precoursor Error | ppm error of the precursor |

| | |
|---|---|
| Precoursor Absolute Error | absolute value of the precursor error |
| PrecoursorAbsoluteErrorRelative | the absolute error/max precursor error |
| 1-ErrorRelative | 1-PrecoursorAbsoluteErrorRelative |
| AverageMS2Error | The average error of all ms2 peak matches |
| MeanSquareError | MSE of the MS2 peak matches |
| MeanSquareRootError | MSRE of the MS2 peak matches |
| AverageRelativeMS2Error | mean of peak error/max error |
| Average1-RelativeMS2Error | mean of (1-peak error/max error) |
| AverageMS2ErrorPeptide1 | Average error of all linear fragment matches for peptide1 |
| AverageMS2ErrorPeptide2 | Average error of all linear fragment matches for peptide2 |
| AverageMS2ErrorCrossLinked | Average error of all cross-linked fragments |

#### MS2 Fragment Charges

| *Field* | *Description* |
|---|---|
| MaxCharge | maximal observed charge |
| AverageCharge | average observed charge |
| RelativeMaxCharge | MaxCharge/Precursor Charge |
| RelativeAverageCharge | AVerageCharge/Precursor Charge |

#### Fragment occurrence

| *Score* | *Description* | *Name as Applied To* | | |
|---|---|---|---|---|
| | | *Whole match* | *for Peptide1* | *for Peptide2* |
| total fragment matches | Number of fragment to peak matches | total fragment matches | peptide1 total fragment matches | peptide2 total fragment matches |
| lossy fragment matches | Number of neutral loss fragment to peak matches | lossy fragment matches | peptide1 lossy fragment matches | peptide2 lossy fragment matches |
| matched | Number of fragments matched (each unique b/y ion is only counted ones) | fragment matched | peptide1 matched | peptide2 matched |
| coverage | found fragmentation site/maximum fragmentation sites | fragment coverage | peptide1 coverage | peptide2 coverage |
| non lossy matched | Non neutral loss | fragment non | peptide1 non lossy | peptide2 non |

| | fragments matched | lossy matched | matched | lossy matched |
|---|---|---|---|---|
| non lossy coverage | Non neutral loss coverage | fragment non lossy coverage | peptide1 non lossy coverage | peptide2 non lossy coverage |
| lossy matched | Neutral loss fragments matched | fragment lossy matched | peptide1 lossy matched | peptide2 lossy matched |
| lossy coverage | Neutral loss fragment coverage | fragment lossy coverage | peptide1 lossy coverage | peptide2 lossy coverage |
| matched conservative | same as non-lossy coverage but if a fragment is observed in three distinct neutral loss states then it is a also counted as non-lossy | fragment matched conservative | peptide1 matched conservative | peptide2 matched conservative |
| conservative coverage | as fragment matched conservative but as coverage | fragment conservative coverage | peptide1 conservative coverage | peptide2 conservative coverage |
| unique matched | same as "matched" but for each peak only one fragment match is considered | fragment unique matched | peptide1 unique matched | peptide2 unique matched |
| unique matched non lossy | same as "matched non-lossy" but for each peak only one fragment match is considered | fragment unique matched non lossy | peptide1 unique matched non lossy | peptide2 unique matched non lossy |
| unique matched non lossy coverage | same as "matched non-lossy-coverage" but for each peak only one fragment match is considered | fragment unique matched non lossy coverage | peptide1 unique matched non lossy coverage | peptide2 unique matched non lossy coverage |
| unique matched lossy | same as "matched lossy" but for each peak only one fragment match is considered | fragment unique matched lossy | peptide1 unique matched lossy | peptide2 unique matched lossy |
| unique matched lossy coverage | same as "matched lossy coverage" but for each peak only one fragment match is considered | fragment unique matched lossy coverage | peptide1 unique matched lossy coverage | peptide2 unique matched lossy coverage |
| unique matched conservative | same as "matched conservative" but for each peak only one fragment match is considered | fragment unique matched conservative | peptide1 unique matched conservative | peptide2 unique matched conservative |
| unique matched conservative coverage | same as "matched coverage" but for each peak only one fragment match is considered | fragment unique matched conservative coverage | peptide1 unique matched conservative coverage | peptide2 unique matched conservative coverage |
| multimatched% | how many of the matched fragments are matched in more then one charge state | fragment multimatched% | peptide1 multimatched% | peptide2 multimatched% |

| | | | | |
|---|---|---|---|---|
| sequencetag coverage% | how much of the peptides sequence is explained by consecutive fragments | fragment sequencetag coverage% | peptide1 sequencetag coverage% | peptide2 sequencetag coverage% |

#### Search-Database derived Scores

| *Field* | *Description* |
|---|---|
| FragmentLibraryScore | how specific are all peak taht are matched to the given peptide versus all other peptides |
| FragmentLibraryScoreLog | ln(FragmentLibraryScore) |

#### Spectrum coverage

| | |
|---|---|
| spectrum intensity coverage | how much of the total intensity of the spectrum is explained |
| spectra intensity nonlossy coverage | how much of the total intensity of the spectrum is explained by non-neutral loss fragments |
| spectra isotop% | how much of the spectrum intensity is contained in isotope clusters |
| spectra matched isotop% | how much of the spectrum isotope cluster are actually matched to something |
| spectra matched single% | how much of the single peaks are matched |
| spectra top10 matched% | how much of the top n peaks are explained |
| spectra top20 matched% | |
| spectra top40 matched% | |
| spectra top100 matched% | |
| spectrum peaks coverage | how much of the total number of peaks are matched |
| SpectraCoverageConservative | how much of the spectrum is explained either by fragments found as bassic (non-neutral loss) fragments + fragments that are not found as non-neutral loss but at least with three distinct neutral loss fragments |

#### Preliminary Combinations of Sub-Scores

| | |
|---|---|
| Pep1Score | some weighted combination of sub-scores pertaining to peptide1 |
| Pep2Score | some weighted combination of sub-scores pertaining to peptide2 |
| spectrum quality score | some weighted combination of sub-scores pertaining to the spectrum |

#### Other Sub-Scores

| | |
|---|---|
| Autovalidation | a ml model defining a match as being validated - this is the base of the "validated" column but is there filtered by additional criteria |

#### Legacy Sub-Scores

Subscores that are still present but should no longer be considered

| |
|---|
| FragmentLibraryScoreExponential |
| BS3ReporterIonScore |
| Crosslinked |
| Modified |
| Containing |
| J48ModeledManual001 |
| RandomTreeModeledManual |
| AllScore |
| AllScoreLib |
| MatchScore |
| NormScore |

#### Final Score

the sub-scores get normalized, weighted and averaged to calculated the final score for the match
This is the score in the last column of the CSV-file

| | |
|---|---|
| **match score** | **main score of the PSM** |
